# Identification of a long non-coding RNA regulator of liver carcinoma cell survival

**DOI:** 10.1101/2020.07.01.181693

**Authors:** Yulia Rybakova, John T. Gonzalez, Roman Bogorad, Vikash P. Chauhan, Yize L. Dong, Charles A. Whittaker, Victor Koteliansky, Daniel G. Anderson

## Abstract

Genomic studies have significantly improved our understanding of hepatocellular carcinoma (HCC) biology and have led to the discovery of multiple protein-coding genes driving hepatocarcinogenesis. In addition, these studies have identified thousands of new non-coding transcripts deregulated in HCC. We hypothesize that some of these transcripts may be involved in disease progression. Long non-coding RNAs are a large class of non-coding transcripts which participate in the regulation of virtually all cellular functions. However, a majority of lncRNAs remain dramatically understudied. Here, we applied a pooled shRNA-based screen to identify lncRNAs essential for HCC cell survival. We validated our screening results using RNAi, CRISPRi, and antisense oligonucleotides. We found a lncRNA, termed ASTILCS, that is critical for HCC cell growth and is overexpressed in tumors from HCC patients. We demonstrated that HCC cell death upon ASTILCS knockdown is associated with apoptosis induction and downregulation of a neighboring gene, Protein Tyrosine Kinase 2 (PTK2), a crucial protein for HCC cell survival. Taken together, our study describes a new, non-coding RNA regulator of HCC.

## INTRODUCTION

Liver cancer is one of the leading causes of cancer mortality worldwide, accounting for more than 700,000 deaths per year [1]–[3]. Hepatocellular carcinoma (HCC) is the most frequent subtype of liver cancer. Despite recent progress in HCC treatment it remains one of the deadliest types of cancer [3], [4]. Notably, the incidence of HCC has been increasing in recent decades, making HCC one of the fastest-growing causes of death worldwide [5], [6]. This poor prognosis underlines the need for new effective therapies. Better understanding of the molecular mechanisms regulating HCC progression may yield new potential drug targets.

A meta-analysis of human HCC datasets revealed 935 genes for which expression was significantly dysregulated in HCC samples compared to healthy tissues [7]. Further Gene Ontology analysis of these genes identified several gene networks associated with HCC progression. Among them were upregulation of cell proliferation, downregulation of apoptosis, loss of hepatocyte differentiation, immunosuppression, and activation of proteins acting at an epigenetic level [7]. Comprehensive genomic profiling of patient HCC samples and their comparison with healthy tissues have helped uncover molecular changes promoting the above phenotypic features of HCC [8]–[15]. Among them, mutations leading to activation of the WNT signaling pathway were most common [9]–[14], [16], implicating the WNT pathway as a major driver of hepatocarcinogenesis [17], [18]. Moreover, activation of the WNT pathway is associated with an immunosuppressive microenvironment, another hallmark of HCC progression [8], [15], which emphasizes the role of WNT pathway activity in HCC progression. Other common mutations affected the TERT promoter, TP53, genes regulating cell cycle, PI3K-AKT-mTOR signaling and cell differentiation [9]–[14]. Notably, up to 50% of clinical HCC samples reported in different studies have a mutation in chromatin modifiers [9]–[13], indicating the importance of epigenetic regulation in HCC development.

Besides shedding light on the roles of protein-coding genes, integrative genomic studies have revealed that the majority (>70%) of transcribed sequences in the human genome participate in cell function regulation without producing a protein [19], [20]. Long non-coding RNAs (lncRNAs) are defined as non-coding transcripts longer than 200 nucleotides and represent a large class of non-coding elements, comprising more than 50,000 annotated transcripts to date [21]–[25]. Pertinently, hundreds of lncRNAs are recurrently deregulated in HCC, suggesting potential roles in hepatocarcinogenesis. Co-expression network analysis determined that these lncRNAs were associated with cell proliferation, metastasis, immune response, and liver metabolism – hallmarks of HCC progression [26]–[29]. While the pathogenic roles of some of these lncRNAs (e.g. HULC, H19, HOTAIR, HOTTIP, DANCR) have already been described [30]– [32], a plurality of lncRNA transcripts remain largely uncharacterized. Discovery of novel lncRNAs and their intracellular functions promises to expand our knowledge of HCC cellular physiology and may provide the basis for new therapeutic modalities.

Currently, lncRNA functions cannot easily be predicted based on their sequence. Instead, subcellular localization, transcript abundance, and functional genomic screens can help to efficiently narrow down possible lncRNA biological roles and molecular functions [33]–[37]. For instance, lncRNAs located mainly in the nucleus typically function as transcription regulators of local genes (*in cis*) or distant genes (*in trans*) [36]. Cytoplasmic lncRNAs are more likely to regulate protein production, formation of post-translational modifications, and sequestration of miRNAs or RNA-binding proteins [37]. Transcript abundance can provide another hint about lncRNA function. For example, low-abundance transcripts tend to function in *cis* because their low concentration makes diffusion a barrier to activity at long distances from the transcription site. Abundant lncRNAs, on the other hand, can achieve high concentrations at multiple target regions, including those outside of the nucleus and therefore often function in *trans* [33]. Finally, pooled functional genetic screens are a powerful tool allowing for parallel perturbation of multiple genes to select for those that are critical for a phenotype or function [38]–[40]. Recently, genome-wide screens have made it possible to identify lncRNAs involved in a wide variety of cellular functions including cell proliferation, drug resistance, autophagy, tissue homeostasis, and cell differentiation [41]–[46].

RNA interference (RNAi) is an effective method for transient silencing of gene expression and therefore is an instrument for loss-of-function genetic screens [38], [40]. Previously, it was reported that RNAi-mediated gene silencing is restricted to the cytoplasm, limiting targeting of nuclear transcripts. However, recent studies suggest RNAi presence and activity in the mammalian nucleus as well, although with less efficiency [47]–[50]. Clustered regularly interspaced short palindromic repeat interference (CRISPRi) is another potent technique for lncRNA silencing [39], [51], [52]. However, using CRISPRi to regulate a lncRNA overlapping with other transcripts might contribute to the expression of that transcript, confounding data interpretation [53]. Given promoters of most lncRNAs are poorly annotated and lncRNAs often overlap with protein-coding genes (or their promoters/enhancers), in our screen, we chose to perturb lncRNA at the RNA level. We performed an shRNA-based pooled screen to identify lncRNAs essential for the survival of the human HCC cell line HUH7. Based on the lncRNA expression profile of these cells, we designed a lentiviral shRNA library targeting all identified lncRNAs. Using this library, we performed a loss-of-function genetic screen and found that lncRNA ENST00000501440.1 is critical for HUH7 cell growth. We named this lncRNA ASTILCS (**A**nti**S**ense **T**ranscript **I**mportant for **L**iver **C**arcinoma **S**urvival). Importantly, in patient data, ASTILCS is significantly overexpressed in HCC compared to normal tissues. Further, using gene manipulation techniques, we demonstrate that ASTILCS knockdown results in apoptosis induction and HCC cell death. Finally, we show that ASTILCS knockdown correlates with downregulation of a neighboring gene expressing Protein Tyrosine Kinase 2 (PTK2), the silencing of which results in HCC cell death.

## RESULTS

### Pooled RNAi-based screen identifies lncRNAs potentially essential for HCC cell survival

To design the shRNA library, we performed transcriptome analysis in HUH7 HCC cell line and identified 1618 non-coding RNA transcripts longer than 200 base pairs and expressed at a level higher than 5 FPKM (Supplemental Fig. 1, Supplemental Table 1). Next, we constructed a library of 7873 shRNA vectors to knockdown the identified lncRNAs based on RNAi and applied on HUH7 cells (Fig. 1A). Each lncRNA was targeted by 4-5 shRNAs to account for shRNA off-target effects. To identify lncRNAs important for HUH7 cell survival, shRNAs present in the final population were compared to shRNA representation in the input library. A lncRNAs was considered a candidate when at least two of its corresponding shRNAs were underrepresented in the final population with log_2_(fold change compared to control) ≥ 1 or by at least 3 shRNAs with log_2_(fold change compared to control) ≥ 0.75 (Fig. 1B). With these constraints, we identified seven lncRNA candidates for further validation (ENST00000429829, ENST00000510145, ENST00000457084, ENST00000501440.1, ENST00000366097.2, ENST00000518090 and ENST00000421703.5). To the best of our knowledge only ENST00000429829 and ENST00000510145 very previously characterized [54]–[60].

**Figure 1.**
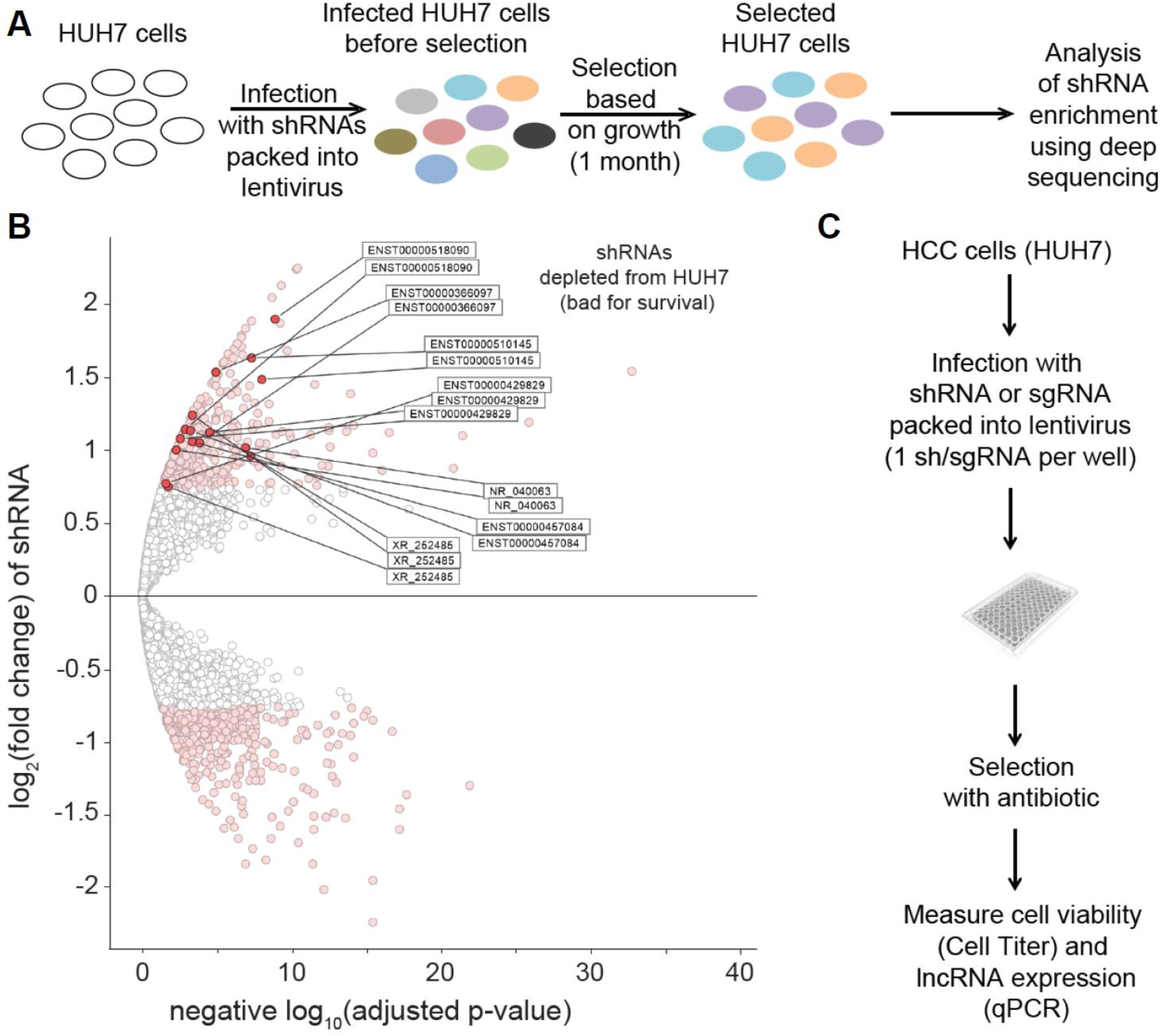
Experimental design and selection strategy for the identification of lncRNAs essential for HUH7 HCC cell survival. **A.** Schematic workflow of the survival-based pooled shRNA library screen in HUH7 cells. shRNAs were designed to target lncRNAs identified in the cell line. **B.** Volcano plot of the differentially expressed shRNAs in the final population of HUH7 cells. The x-axis indicates the adjusted p values plotted in −log10. The y-axis indicates the log2(fold change) in gene expression, which was defined as the ratio of normalized gene expression in the input library over the final HUH7 population. Light red dots represent shRNAs with log2FC≥0.75 and adjusted p-value≤0.05. Dark red dots represent shRNAs of lncRNAs for which at least 2 shRNAs have log2FC≥ 1 and adjusted p-value≤0.05 or lncRNAs for which at least 3 shRNAs have log2FC≥0.75 and adj p-val≤0.05. **C.** Schematic workflow of arrayed shRNA and sgRNA screens used for validation of lncRNAs identified in B.

Both of these lncRNAs have been identified in the context of cancer. Although their mechanisms are the focus of active discussion, their presence among our screen hits supports the likelihood that the rest of the transcripts are also involved in HCC survival and biology. ENST00000429829 is one of the multiple transcripts of gene ENSG00000229807, also known as XIST. In addition to its established role as the master regulator of X chromosome inactivation [61], XIST has been reported to participate in progression of a variety of cancers, including HCC [54]– [58], [62]–[64]. However, the results of these studies are controversial [54]–[58]. ENST00000510145 is one of nine transcripts of gene ENSG00000250682, also known as LINC00491 or BC008363. This gene was found to be upregulated in a TCGA colon adenocarcinoma dataset and was associated with lower patient survival, implying ENSG00000250682 importance for colorectal cancer progression [59]. Conversely, in pancreatic ductal adenocarcinoma patients LINC00491 expression was significantly lower compared to the control group and was associated with better survival rates [60].

### Validation of the screen results identifies lncRNA ASTILCS as a new regulator of HCC cell survival

To validate the screening results, we individually expressed the five library shRNAs for each of the seven candidate lncRNAs (Supplemental Table 2) and repeated the screen in an arrayed format (Fig. 1C). Those lncRNAs for which at least two corresponding shRNAs reduced cell survival by more than 50% compared to the control shRNAs were selected for further analysis; these were ENST00000501440.1, ENST00000366097.2, ENST00000518090, and ENST00000421703.5 (Fig. 2A).

**Figure 2.**
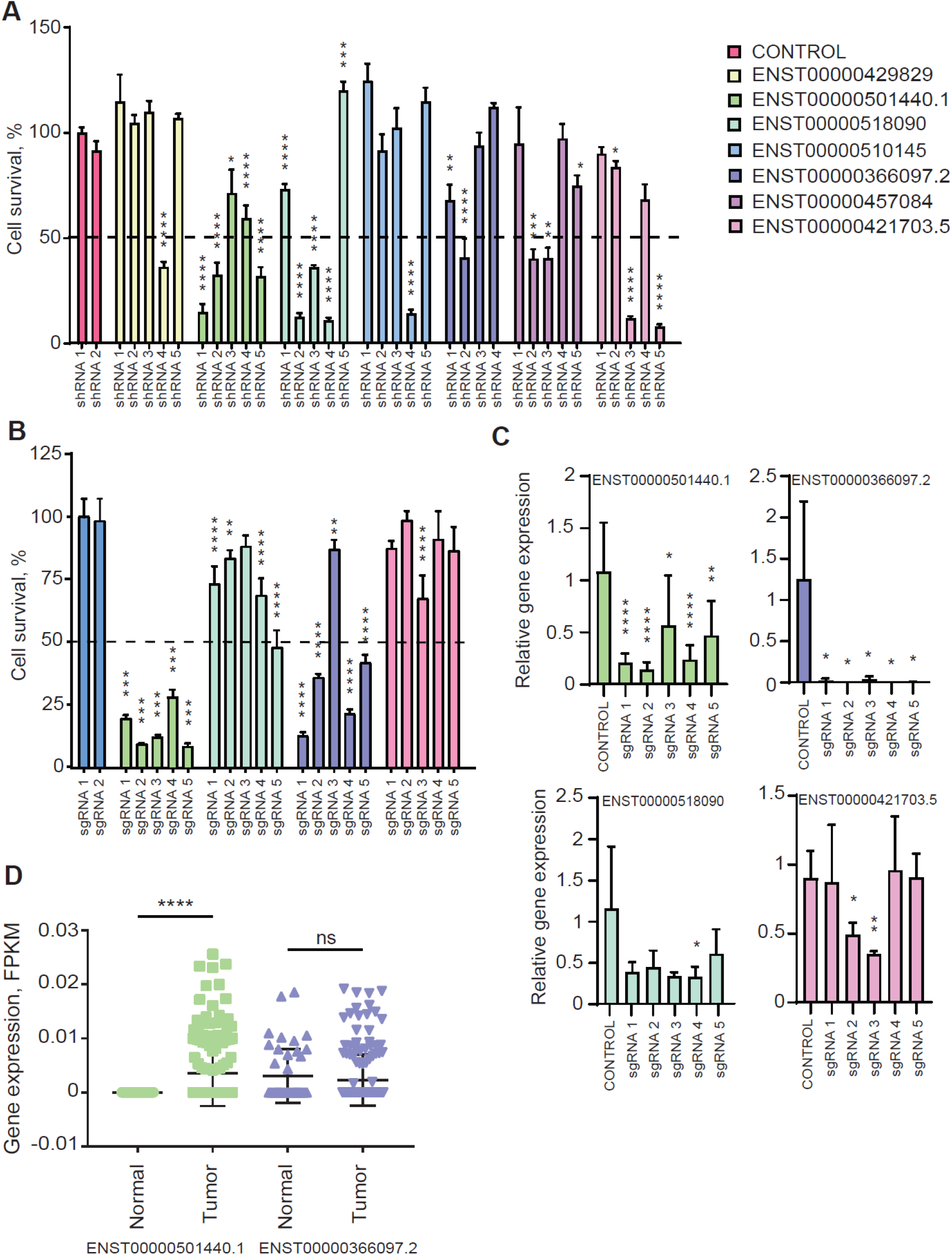
Validation of the screen results identifies lncRNA ASTILCS a new regulator of HCC cell survival. **A.** HUH7 cell survival upon shRNA-mediated knockdown of candidate lncRNAs (compared to shRNA1), n≥3. **B.** HUH7 cell survival upon CRISPRi-mediated knockdown of candidate lncRNAs from A. Compared to control sgRNA1, n≥3. **C.** LncRNA expression in HUH7 cells transduced with sgRNA-dCas9-KRAB targeting one of the candidate lncRNAs, n≥4. **D.** ENST00000501440.1 (ASTILCS) and ENST00000366097.2 expression in HCC vs adjacent tissue (TCGA-LIHC-rnaexp dataset#), n≥45. All values are mean ± SD, **** p < 0.0001; *** p < 0.001; **p < 0.01; *p < 0.05.

RNAi-based gene silencing is associated with a few pitfalls, particularly off-target activity and variability in knockdown efficiency [65], [66]. We therefore further validated the candidate lncRNAs using an arrayed screen based on CRISPRi (Fig. 1C). To do so, we designed five sgRNAs (Supplemental Table 3) to allow targeting of each candidate lncRNA by CRISPRi (Fig. 2B, C). Among the four studied lncRNAs, CRISPRi-mediated knockdown of only ENST00000501440.1 and ENST00000366097.2 resulted in substantially decreased survival for HUH7 HCC cells (Fig. 2B, C). Specifically, 5/5 sgRNAs targeting lncRNA ENST00000501440.1 decreased HCC cell survival by more than 70% and 4/5 sgRNAs targeting lncRNA ENST00000366097.2 resulted in more than 50% HUH7 cell death. In contrast, knockdown of ENST00000518090 was not associated with a notable decrease in HCC cell survival and only 2/5 sgRNAs designed to target ENST00000421703.5 induced partial lncRNA knockdown with mild effects on HCC cell survival. Based on these results we concluded that ENST00000501440.1 and ENST00000366097.2 expression is critical for HCC cell survival. ENST00000501440.1 is the only transcript of ENSG00000244998 gene. It is a 1380 bp long antisense transcript comprised of 2 exons. ENST00000366097.2 is one of two transcripts of ENSG00000203266 gene. It is a 770 bp long intergenic lncRNA consisting of 3 exons. Both transcripts are predicted to have low coding potential and are not conserved in chimpanzee or mouse [67]. Thus, we identified two novel lncRNA genes which expression is potentially important for HCC cell survival.

To determine whether these two lncRNAs are HCC specific or are present in healthy liver tissues, we examined ENST00000501440.1 and ENST00000366097.2 expression in tissue samples from patients with HCC using a dataset from The Cancer Genome Atlas (TCGA-LIHC-rnaexp, downloaded from The Atlas of NcRNA in Cancer (TANRIC) [68]). We found that ENST00000501440.1 expression was significantly higher in liver cancer samples compared to the adjacent tissue (Fig. 2D; p<0.0001). Yet, lncRNA expression was not associated with patient survival [68]. These data suggest that only ENST00000501440.1 expression is critical for the survival of tumor cells. Because only ENST00000501440.1 expression is differentially expressed in cancer cells, we selected it for further analysis. Through the rest of the publication, we refer to this lncRNA by the name of ASTILCS (**A**nti**S**ense **T**ranscript **I**mportant for **L**iver **C**arcinoma **S**urvival).

A closer look into the ASTILCS locus revealed that ASTILCS is an antisense sequence to the protein-coding gene Protein Tyrosine Phosphatase Type IVA 3 (PTP4A3) (Supplemental Fig. 2). PTP4A3 is known to be important for cell proliferation; its knockdown has been shown to decrease survival in multiple types of cells [69]–[73]. Because sgRNAs targeting ASTILCS bind PTP4A3 between 512 and 611 bp away form the transcription start site, there is a possibility that the sgRNA-dCas9-KRAB complex hinders PTP4A3 expression, resulting in HCC cell death independently of ASTILCS. Indeed, expression analysis of the sgRNA treated cells revealed deep knockdown of PTP4A3 (Supplemental Fig. 3). To add orthogonal evidence of ASTILCS prosurvival effects on HCC cells, we knocked down its expression by transient transfection of antisense oligonucleotides containing locked nucleic acid modifications (LNA) (Supplemental Table 4). LNAs bind with high affinity to complementary RNA sequences forming DNA•RNA hybrids, which are recognized and cleaved by RNAse H1, resulting in gene knockdown [74]–[76]. We observed a reduction in HUH7 HCC cell survival upon treatment with the LNAs (Fig. 3A), which was associated with ASTILCS knockdown (Fig. 3B). We noticed that, despite a decrease in cell survival in LNA2-treated samples, ASTILCS RNA levels in these samples were not affected. These findings may be explained by previous reports demonstrating that antisense oligonucleotide hybridization with RNA can affect its function without inducing degradation [77], [78]. Thus, LNA2 binding to ASTILCS might perturb its function via steric blocking of lncRNA secondary structure formation or interaction with molecules important for the lncRNA signaling [79]–[81]. To further corroborate whether ASTILCS expression is critical for HCC cell survival, we measured its expression in HUH7 HCC cells transfected with the 3 most efficient shRNAs from the library (Supplemental Fig. 4) and observed dosage-dependent decrease in HCC cell survival (Fig. 3C and Fig. 2A). These findings substantiate that ASTILCS regulates HCC cell survival and its specific knockdown leads to HCC cell death independently of its reciprocal sense coding gene, PTP4A3.

**Figure 3.**
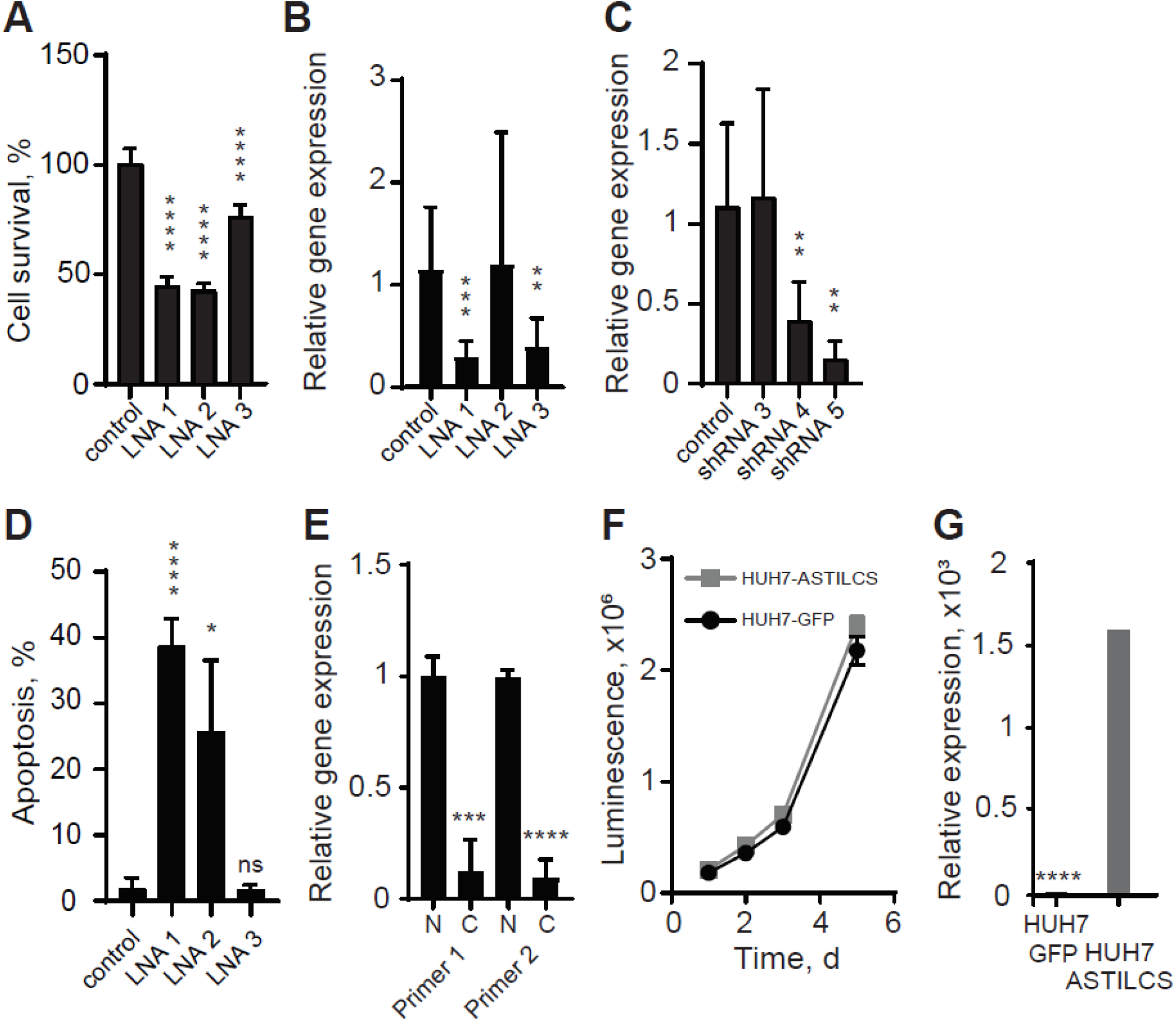
ASTILCS expression is essential for liver carcinoma cell survival. **A.** HUH7 cell survival 48h after transfection with LNAs targeting ASTILCS, n≥6. **B.** ASTILCS expression in HUH7 cells transfected with LNAs targeting ASTILCS, n≥5. **C.** ASTICLS expression in HUH7 cells transduced with shRNAs targeting ASTILCS, n≥5. **D.** Apoptosis in HUH7 cells treated with LNAs targeting ASTILCS, n=3. **E.** LncRNA expression in nucleus and cytoplasm, n≥8. **F.** Growth curve for HUH7 cells transfected with GFP and ASTILCS. **G.** ASTILCS expression in HUH7 cells transduced with ASTILCS-TRC209, n=3. All values are mean ± SD, **** - p < 0.0001; *** - p < 0.001; ** - p < 0.01; * - p < 0.05; ns. - P > 0.05.

### LncRNA ASTILCS knockdown in HCC cells results in apoptosis induction

To understand the molecular mechanism of the effects of ASTILCS on HCC cell survival, we studied whether ASTILCS knockdown affects HUH7 HCC cell apoptosis. To that end, we performed a TUNEL assay to assess apoptosis. We found that transformation with shRNA expressing plasmids or treatment with LNAs led to a dose-dependent increase in the number of apoptotic cells (Fig. 3D and Supplemental Figure 5). Differences in the apoptotic cell number between shRNA and LNA treated samples were likely due to experimental constraints in the knockdown techniques. Apoptosis levels in LNA-treated samples were measured 24 h after the treatment, while in shRNA-treated samples apoptosis could only be measured 4 days after transduction, providing time for compensation mechanisms to occur. Moreover, cell media in shRNA-treated samples had to be changed to remove the lentiviral particles and add selective agent, which could also result in partial removal of poorly attached apoptotic cells. From our findings we conclude that ASTILCS knockdown results in the induction of apoptosis and a subsequent decrease in HUH7 cell survival.

### ASTILCS is a nuclear antisense transcript which functions in *cis*

As subcellular localization can hint towards the molecular mechanism of a lncRNA, we measured ASTILCS transcript levels in nuclear and cytoplasmic extracts and found ASTILCS RNA to be strongly enriched in the nucleus (Fig. 3E). These results are in line with the relatively low expression level of ASTILCS in HUH7 cells (~23.5 FPKM, Supplemental Figure 5), a common feature of nuclear transcripts. Further, to classify the mechanism by which ASTILCS knockdown decreases HCC cell survival, we determined whether ASTILCS functions in *cis* or *trans*. To do so, we overexpressed cDNA encoding ASTILCS from a randomly integrated lentivirus and assessed cell proliferation as the population doubling time (Td). We found that the Td of cells overexpressing ASTILCS (1.13±0.07 days) was similar to the Td of control cells expressing green fluorescent protein (GFP) from the same vector (1.13±0.03 days, p=0.17) (Fig. 3F, G and Supplemental Table 5). Because we did not observe any gain in survival for cells overexpressing ASTILCS, we concluded that ASTILCS is not likely to act in *trans* and that its effects on HCC cell survival are probably associated with *cis* functions.

### ASTILCS silencing is associated with downregulation of neighboring gene PTK2 essential for HCC cell survival

The effects of low abundance nuclear cis-acting lncRNAs occur typically in the loci from which they are transcribed. Those effects can be mediated by: 1) the lncRNA transcripts themselves; 2) the act of lncRNA transcription; or 3) the regulatory DNA elements within the lncRNA locus [82], [83]. To determine whether the investigated phenotype might result from ASTILCS transcript effects on local gene expression, we examined the impact of ASTILCS knockdown on the expression of all genes within 1 Mb of the target site (Fig. 4A). Analysis of the HUH7 HCC cell transcriptome revealed that G Protein-Coupled Receptor 20 (GPR20) and Maestro Heat Like Repeat Family Member 5 (MROH5) are not expressed in HUH7 cells (Supplemental Table 6), so they were removed from consideration. We found that LNA-induced ASTILCS knockdown led to a change in expression of all studied genes in the locus (Fig. 4B). Only downregulation of Solute Carrier Family 45 Member 4 (SLC45A4), Protein Tyrosine Kinase 2 (PTK2), DENN Domain Containing 3 (DENND3) and Trafficking Protein Particle Complex 9 (TRAPPC9) led to a dose-dependent decrease in both ASTILCS expression and HCC cell survival (Fig. 4B, see also Fig. 3A and B). In contrast, shRNA-mediated knockdown of ASTILCS was associated with downregulation of SLC45A, PTK2, and Chromatin accessibility complex protein 1 (CHRAC1) (Fig. 4C, see also Fig. 2A and 3C). Genes that were inconsistent across ASTILCS knockdown approaches were considered to be results of indirect or off-target effects. Because only SLC45A and PTK2 expression was affected in the same manner by both shRNAs and LNAs, we inferred that ASTILCS knockdown potentially induces HCC cell death via downregulation of one or both of these genes. Changes in expression of other genes might be a result of indirect effects of ASTICLS downregulation or simply off-target effects of the LNAs and shRNAs.

**Figure 4.**
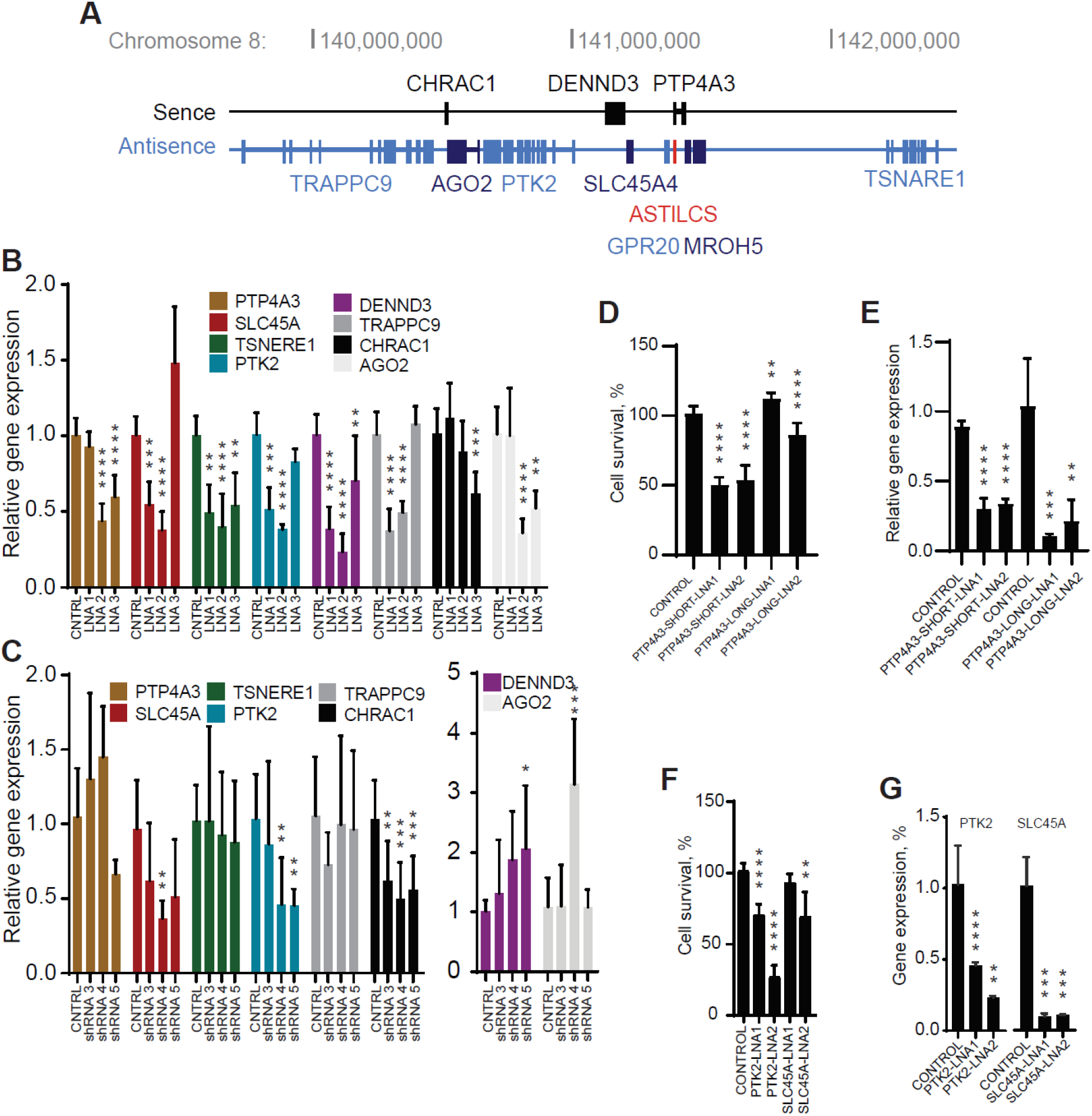
Knockdown of ASTILCS results in dose-dependent downregulation of neighboring genes. **A.** Genomic locus of ASTILCS. Expression of ASTILCS neighboring genes in HUH7 cells upon LNA-mediated (n≥8) (**B**) or shRNA-mediated (n≥5) (**C**) knockdown of ASTILCS. Cell survival (n≥8) **(D)** and gene expression (n≥5) **(E)** upon LNA-mediated silencing of PTP4A3 isoforms. Cell survival (n≥9) **(F)** and gene expression (n≥5) **(G)** upon LNA-mediated silencing of ASTILCS neighboring genes PTK2 or SCL45A, n≥9. All values are mean ± SD, ****p < 0.0001; ***p < 0.001; **p < 0.01; *p < 0.05.

LncRNAs located antisense to protein coding genes are often found to regulate activity of their sense pair in different manners [84], [85]. Surprisingly, even though protein-coding gene PTP4A3 is located antisense to ASTILCS, we did not observe an apparent effect of ASTILCS knockdown on PTP4A3 expression (Fig. 4B, C). This indicates that the ASTILCS transcript itself does not affect the expression of PTP4A3. Next, we studied whether PTP4A3 knockdown can affect HUH7 HCC cell survival. PTP4A3 produces six transcripts (T1-6), three longer (T3-5) than others (T1,2,6) (Supplemental Fig. 6); the sequence of only the long transcripts overlaps with ASTILCS. We designed LNAs targeting long isoforms of PTP4A3 (T3-5) – PTP-LONG-LNA and LNAs targeting two (T1,2) out of three short isoforms of PTP4A3. We could not design an LNA targeting only isoform T6 because it completely overlaps with the long isoforms. Interestingly, we found that knockdown of only the short PTP4A3 isoforms led to a dose-dependent decrease in HCC cell survival (Fig. 4D, E). We also analyzed whether knockdown of the long PTP4A3 isoforms can affect the expression of the short isoforms. With this mechanism in mind, we measured the expression of the short isoforms in HUH7 HCC cells treated with LNAs targeting long isoforms and observed no difference in expression (Supplemental Fig. 7). Because ASTILCS overlaps only with the long PTP4A3 isoforms and their knockdown does not affect the expression of their short, survival modulating counterparts, we conclude that ASTILCS silencing does not lead to a decrease in cell survival via downregulation of PTP4A3 transcripts.

Finally, we studied whether knockdown of SLC45A and PTK2 itself can decrease HCC cell survival. PTK2 has been previously shown to affect HCC cell survival, with PTK2 silencing in HepG2 and HUH6 HCC cells lines reducing cell growth and inducing apoptosis [86]. Meanwhile, SLC45A4 has not been reported to affect cell survival. We treated HUH7 HCC cells with LNAs targeting SLC45A or PTK2 and measured cell survival. We observed that only knockdown of PTK2 was associated with a decrease in HCC cell survival (Fig. 4F, G). Based on our results we conclude that ASTILCS knockdown might decrease HUH7 cell survival and induce apoptosis via downregulation of PTK2.

## DISCUSSION

Despite recent progress in HCC management, it remains the second deadliest cancer type with a 5-year relative patient survival rate of only 18% [1]–[4]. A better understanding of HCC biology informs the development of more efficient treatment strategies. An increasing number of studies suggests a vital role for lncRNAs in HCC progression [32], [87], [88]. However, their functions in HCC biology remain largely unexplored. To address that problem, in our study, we performed an shRNA-based pooled functional genetic screen to find lncRNAs that play crucial roles in HCC cell progression. Applying stringent filtering criteria and three-step validation we identified lncRNAs ASTILCS to be important for survival of HCC cells. To the best of our knowledge, we provide the first characterization of the lncRNA ASTILCS. Following a framework suggested by Joung et al in [43] we determined that ASTILCS is a nuclear lncRNAs with a local regulatory mechanism. Using gene manipulation techniques, we demonstrated that ASTILCS loss-of-function results in apoptosis and downregulation of the neighboring gene PTK2, suggesting a possible mechanism of ASTILCS antisurvival effect.

PTK2, also known as Focal Adhesion Kinase, is a protein tyrosine kinase that plays an essential role in formation of cell-matrix junctions (focal adhesion), regulation of cell migration, and viability in a variety of cell types [89]–[92]. PTK2 recruitment to focal adhesions triggers PTK2 phosphorylation, creating a docking site for SH2 domain-containing proteins (Grb2, Shc etc), thus, linking PTK2 to the activation of the pro-proliferative and anti-apoptotic RAS pathway. Besides that, under certain cellular stress conditions, PTK can be recruited to the nucleus to facilitate Mdm2-dependent ubiquitination of tumor suppressor protein p53 and downregulate apoptosis [93]. Multiple studies report on the importance of PTK2 for cancer progression [94], [95]. To date a few PTK2 inhibitors have been studied in clinical trials, however, the best observed response was stable disease [96]–[98]. Understanding of mechanisms of PTK2 regulation might help to develop more effective PTK2-targeting therapies. Recently, two independent scientific groups simultaneously demonstrated that PTK2 is essential for HCC formation and growth *in vivo* because of its role in activation of the WNT/b-catenin signaling. PTK2 overexpression stimulated β-actin accumulation in the cell nucleus, thereby enhancing transcription of β-actin target genes and promoting hepatocarcinogenesis. PTK2 silencing, on the other hand, led to increase in apoptosis and a decrease in tumor growth [99], [100]. Thus, PTK2 downregulation by ASTICLS knockdown can be an important factor mediating the mechanism of ASTILCS’ proapoptotic effect in HCC cells.

The molecular mechanisms of ASTILCS increasing PTK2 expression will require further studies. Epigenetic regulation might be one of the possible mechanisms. PTK2 is overexpressed in 30-60% of HCC patients and is associated with a higher metastasis rate and reduced survival. Meanwhile, PTK2 expression in healthy liver tissues is negligible, which underlines the importance of PTK2 expression for HCC progression [101]–[104]. In this study, we found that ASTILCS levels were also significantly increased in HCC samples compared to normal tissues. Interestingly, DNA sequence analysis in HCC patient samples revealed that PTK2 is amplified in only 19-26% of cases and mutated in 2.5% [11], [99], [105]–[107]. Therefore, there should be additional epigenetic mechanisms activating PTK2 expression. Examination of the PTK2 promoter demonstrated that the total methylation level of its CpG islands negatively correlated with PTK2 gene expression. Thus, promoter demethylation might be a mechanism of PTK2 overexpression. Indeed, treatment of HCC cells with a demethylation agent has shown to increase PTK2 mRNA and protein levels [99]. Some lncRNAs are known to affect DNA methylation via direct interaction with DNA methyltransferases (DNMTs) or via indirect recruitment of DNMTs through an intermediate protein [108]. Hence, the aforementioned evidence creates a possibility that ASTILCS can increase PTK2 expression via regulation of its promoter methylation.

In addition to examination of ASTILCS effects on PTK2, we explored its relationship with other neighboring genes. One of them, SLC45A4, is a proton-associated sucrose transporter, for which there are no reports of direct association with cancer or cell survival (PubMed search on 03-24-2020). In this study, we demonstrate for the first time, that ASTILCS knockdown leads to SLC45A4 silencing and that SLC45A4 silencing doesn’t affect cell survival in HCC cells. Surprisingly, we did not observe an obvious correlation between knockdown of antisense lncRNA ASTILCS and expression of its sense protein-coding pair, PTP4A3 gene.

Thus, we inferred that the decrease in HCC cell survival upon ASTILCS knockdown is not likely mediated by changes in PTP4A3 expression. PTP4A3, also known as Phosphatase of Regenerating Liver 3 (PRL-3), is a protein-tyrosine phosphatase implicated in both cell proliferation and invasion in several types of cancer, including HCC [109]–[112]. Despite the importance of PTP4A3 for HCC cell survival, it seems the pro-survival effect of PTP4A3 is not regulated by ASTICLS RNA expression. Yet, this does not exclude the existence of other regulatory mechanisms between ASTILCS and PTP4A3 nor their importance in still undiscovered cell functions. Interestingly, the functional analysis of PTP4A3 transcripts presented here suggests that different transcripts affect cell survival in different ways in HCC cells. For the first time, we report that only knockdown of short PTP4A3 transcripts (T1 and T2) reduces the cell survival, while expression of the long transcripts (T3-T5) has no effect on cell viability. This finding is in concordance with functional duality of PTP4A3, which is reported to regulate both cell survival and metastasis. Given only the expression of short transcripts correlates with cell survival, we can speculate that long transcripts might be involved in cell motility and invasion. This hypothesis requires further exploration.

In summary, we identified and characterized a lncRNA, ASTILCS, which regulates HCC cell survival presumably via activation of PTK2 expression and induction of apoptosis. In addition, we unveiled the effects of ASTILCS neighboring genes, PTK2, SLC45A4 and PTP4A3, on HCC cell survival. These findings provide valuable information about HCC biology and can advance the development of future HCC treatments.

## ACKNOWLEDGMENTS

This work was supported by the MIT Skoltech Initiative, Defense Advanced Research Projects Agency (W32P4Q-13-1-001), S. Leslie Misrock Frontier Research Fund for Cancer Nanotechnology, Translate Bio (Lexington, MA), and Koch Institute Support (core) Grant P30-CA14051 from National Cancer Institute. We thank the Koch Institute Swanson Biotechnology Center for technical support, specifically bioinformatics & computing, flow cytometry, genomics (BioMicro Center) and high throughput screening. We also thank Prof. Philip Sharp (MIT) and Prof. Timofei Zatsepin (Skoltech) for valuable discussions of the project, and Dr. Elena Smekalova and Dr. Mikhail Nesterchuk (MIT) for technical support at different stages of the project.

## AUTHOR CONTRIBUTIONS

Y.R. and R.B. designed the study; R.B. performed RNA sequencing and managed RNAi library design; Y.R. executed the rest of the experiments, analyzed data and wrote the manuscript; J.G. assisted with the experiments and edited the manuscript; Y.D. assisted with the experiments; V.C. contributed to experimental design and execution, and edited the manuscript; C.V. analyzed next-generation sequencing data; V.K. and D.A. supervised the study and edited the manuscript.

## MATERIALS AND METHODS

### Cell culture

Human hepatocellular carcinoma HUH7 cell line was a gift from Dr. Jay Horton (UT Southwestern Medical Center). HUH7 and HEK293ft cell lines were grown in Dulbecco’s modified Eagle’s medium with L-glutamine (DMEM, Gibco™) supplemented with 4.5 mg/ml glucose, 50 ug/ml gentamicin sulfate (Sigma), 25 mM HEPES (Gibco™) and 10% heat-inactivated fetal bovine serum (FBS, Gibco™). All cells were cultured at 37°C, 5% CO_2_. When the cells reached a 70-80% monolayer, they were detached from the flask using 0.25% Trypsin-EDTA solution and split 1:10. Concentrations for selection agents were determined using killing curve: 2.5 ug/ml puromycin (Sigma), 0.75 mg/ml G-480 (Sigma).

### RNA sequencing and data analysis

Samples were prepared using strand-specific Ribo-Zero kit and RNA sequencing was performed by MIT BioMicro Center (https://openwetware.org/wiki/BioMicroCenter:Software#BMC-BCC_Pipeline). Reads were aligned to transcripts derived from the hg19 assembly and the Ensembl version 68 non-coding RNA annotation (non-coding genes) or the full Ensembl 68 annotation (protein-coding genes) using Bowtie version 1.01 [113] and gene expression was summarized using RSEM version 1.2.3 [114].

### Genome-wide screening

Based on HUH7 RNA sequencing results (Supplemental Fig.1, Supplemental Table 1), we designed a library of 7873 shRNA vectors allowing to do knockdown of the identified 1618 lncRNAs based on RNAi. The library was developed, synthesized and packed into lentivirus by the RNAi Consortium at the Broad Institute [115]. The shRNA sequences were assembled into a pLKO.1 lentiviral backbone (Addgene plasmid #10878), containing a puromycin resistance marker to allow for the antibiotic selection of transduced cells. CMV-VSV-G (Addgene plasmid #8454) and psPAX2 (Addgene plasmid #12260) plasmids were used for lentiviral packaging. The lentiviral library contained four to five shRNAs per target lncRNA and was applied at a low multiplicity of infection (MOI) equal to 0.3. Two days after lentiviral library exposure, infected cells were selected for four days on puromycin. To assess effects of shRNAs on cell survival, the selected cells were cultured for four more weeks maintaining an shRNA representation of 500 (i.e. each shRNA was expressed on average by 500 cells). The input pooled shRNA plasmid library before virus production was also sequenced and used as a control.

### Next generation sequencing

Samples for Illumina sequencing were prepared following “One Step PCR Preparation of Samples for Illumina Sequencing” protocol from The RNAi Consortium (https://portals.broadinstitute.org/gpp/public/resources/protocols). Briefly, gDNA was isolated using the QIAamp DNA Blood Maxi Kit (Qiagen). Illumina adapter sequences with 5-letter barcodes were used to PCR amplify the shRNA-expressing cassette. The samples were multiplexed and sequenced by MIT BioMicroCenter using HiSeq2000 platform. The samples were processed using the BMC/BCC 1.5.2 pipeline (updated on 08/12/2016). Adapter sequence GGAAAGGACGAGGTACC was trimmed from reads using Cutadapt version 1.4.2 [116]. Trimmed reads were then aligned target consisting of the 7873 sequence shRNA library with BWA version 0.7.10 [117]. Mapped reads were summarized and parsed using SAMtools version 1.3 [118] and custom Perl scripts. The resulting count table was tested for differential representation using DESeq2 version 1.10.1 [119] running under R version 3.2.3. Differential expression data was visualized using Tibco Spotfire Analyst version 7.11.1.

### Molecular cloning

shRNAs from the library (Supplemental Table 2) were annealed and cloned into a pLKO.1_neo plasmid (a gift from Sheila Stewart; Addgene plasmid # 13425; http://n2t.net/addgene:13425; RRID:Addgene_13425) using a protocol from [120]. Two shRNAs designed to target mCherry were used as controls. Briefly, oligos were resuspended in water to a final concentration of 100 uM. 11.25 ul of each oligo (top and bottom) were mixed with 2.5 ul of 10X annealing buffer (1M NaCl, 100 mM Tris-HCl, pH=7.4) and annealed at 95°C using a water bath. The pLKO.1_neo plasmid was digested using AgeI and EcoRI restriction enzymes and purified on 1% agarose gel. Next, oligo mixture was diluted 1:400 in 0.5X annealing buffer and ligated with the digested pLKO.1_neo plasmid using T4 DNA ligase (3 h at RT). 2 ul of the ligation mixture was used to transform 10 ul of One Shot competent Stbl3 E. coli cells (Invitrogen) according to manufacturers’ instructions. Transformed bacteria were plated on LB-agar plates with 100 ug/mL ampicillin and incubated overnight. Individual colonies were picked, inoculated in 3 ml of LB with ampicillin to start miniprep cultures and incubated for 24 h. Miniprep DNA was isolated using QIAGEN Plasmid Mini Kit (Qiagen). shRNA sequences were confirmed by Sanger sequencing (performed by Quintara Biosciences).

sgRNAs (Supplemental Table 3) were designed using the Broad Institute’s GPP sgRNA Designer [121], [122]. Two sgRNAs targeting mouse XIST and blasted against human genome and transcriptome were used as controls. Then, the sgRNAs were assembled into a plasmid expressing dead Cas9 (dCas9, Cas9 without endonuclease activity) fused with a transcription inhibitor, The Krüppel associated box (KRAB) transcriptional repression domain, in a lentiviral backbone containing a puromycin resistance sequence (pLV hU6-sgRNA hUbC-dCas9-KRAB-T2a-Puro, a gift from Charles Gersbach, Addgene plasmid # 71236; http://n2t.net/addgene:71236; RRID:Addgene_71236, [52]) using Golden Gate assembly reaction as described in [123]. 2 ul of the ligation mixture were used to transform 10 ul of NEB Stable Competent E. coli (NEB) according to manufacturers’ instructions. Transformed bacteria were plated on LB-agar plates with 100 ug/mL ampicillin and incubated overnight. Individual colonies were picked, inoculated in 3 ml of LB with ampicillin to start miniprep cultures and incubated for 24 h. Miniprep DNA was isolated using QIAGEN Plasmid Mini Kit (Qiagen). sgRNA sequences were confirmed by Sanger sequencing (performed by Quintara Biosciences).

To create a plasmid expressing ASTILCS, it’s full sequence was used to substitute GFP in TRC209 lentiviral plasmid (PGK-Hygro-EF1a-GFP, gift from the Broad GPP, [43]). The cloning and sequence validation were done by Genscript Biotech.

### Lentivirus production and transduction

For transduction, plasmids were packaged into lentivirus through transfection of the plasmids with a packaging plasmid (psPAX2 was a gift from Didier Trono (Addgene plasmid # 12260; http://n2t.net/addgene:12260; RRID:Addgene_12260)) and an envelope plasmid (CMV-VSV-G was a gift from Bob Weinberg (Addgene plasmid # 8454; http://n2t.net/addgene:8454; RRID:Addgene_8454), [124]) using TransIT-LT1 Transfection Reagent (Mirus Bio). 300 000 HEK293ft cells were plated per well into a 6-well plate and incubated overnight. 0.4 ug PAX2, 0.15 VSV-G and 3.3 ug plasmid of interest were added to 600 ul Opti-MEM and mixed with an equal volume of Opti-MEM containing 4 ul of TranslT-LT1. The mixture was incubated at RT for 20 min and transferred to the well. The volume was brought to 600 ml per well with the culture media and incubated overnight. On the following day, the media was changed. Media with lentiviral particles was collected after 48 hs and snap-frozen in liquid nitrogen. All shRNA/sgRNA plasmids were produced in parallel.

### Arrayed screening

Equal numbers of HUH7 cells (5000) were plated in a 96-well plate and transduced with 5ul of shRNAs or 2 ul of sgRNAs packed into lentiviral particles, so that each well received only one type of shRNA/sgRNA. A plasmid expressing Green Fluorescent Protein (GFP) (pLJM1-EGFP was a gift from David Sabatini (Addgene plasmid # 19319; http://n2t.net/addgene:19319; RRID:Addgene_19319), [125]), but not caring antibiotic resistance marker was also packed into lentiviral particles and used as a positive control for transduction and antibiotic selection. After an overnight incubation the cell media was changed. Two days after the lentiviral transduction, a selection reagent (G-480 or puromycin, respectively) was added to the culture media to select for cells containing the shRNA/sgRNA expressing plasmids. Once the selection was completed (i.e. all non-infected GFP treated cells were dead), cell survival was measured using Cell Titer assay.

### Cell survival assay

HUH7 cell survival was analyzed using the CellTiter-Glo^®^ Luminescent Cell Viability Assay according to the manufacturer’s protocol. Luminescence was measured with the microplate reader Tecan Infinite^®^ 200 PRO.

### Cell proliferation assay

Cells were plated at low density in 96-well plates (2000 cells/well). Cell number analysis using cell titer assay was performed at 1, 2, 3 and 5 days afterwards. For growth curves analysis, doubling time was calculated from the exponential portion of the cell growth curve using the following equation: Td = 0.693t/ln(Nt/N0), where t–time (in days), N0– initial cell number, Nt–cell number on day t.

### Gene expression analysis

For single tube reactions (Fig. 2G) RNA was isolated using Omega Bio-tek’s E.Z.N.A.^®^ Total RNA Kit I isolation kit according to manufacturers’ instructions. Separation and purification of cytoplasmic and nuclear RNA (Fig. 2E) were done using Cytoplasmic and Nuclear RNA Purification Kit (Norgen Biotek Corp.) also following manufacturers’ instructions. Reverse transcription reaction was performed using Applied Biosystems™ High-Capacity RNA-to-cDNA™ Kit and 1 ug of RNA. RNA levels were assessed by qPCR using Power SYBR™ Green PCR Master Mix (Applied Biosystems™). For high-throughput experiments RNA isolation, reverse transcription reaction, and qPCR was performed using Power SYBR™ Green Cells-to-CT™ Kit (Ambion) according to manufacturers’ instructions. TaqMan Fast Advanced Master Mix (Applied Biosystems™) was used with TaqMan primers (Hs01060665_g1 for ACTB and Hs01377184_m1 for ASTILCS). β-actin mRNA was used as a housekeeping control. The RNA levels were first normalized to the level of β-actin and then to an average value of the control group. All SYBR Green primers are listed in Supplemental Table 7.

### LNA transfection

LNAs targeting ASTILCS, SLC45A, PTK2 and PTP4A3 genes were custom-designed using Qiagen’s Antisense LNA GapmeR designer (Supplemental Table 4), and non-targeting LNA (Negative Control A (NCA)) was included as a control. LNAs were resuspended in water to a final concentration of 50 uM. 10 000 HUH7 cells were plated per well in a 96-well plate and incubated overnight. LNAs were formulated with Lipofectamine 2000 (Invitrogen) in Opti-MEM (Gibco™) according to the manufacturer. Each well was treated with 50 ul Opti-MEM containing 20 pmol LNA formulated and 0.25 ul Lipofectamine 2000. Cell survival and gene expression were measured 24h after transfection.

### Apoptosis analysis

was performed using In Situ Cell Death Detection Kit, TMR red (Roche) according to manufacturers’ instructions. Briefly, cells were collected using 0.25% Trypsin-EDTA solution, fixed with 4% paraformaldehyde in PBS at RT for 30 min and permeabilized with 0.1% Triton X-100 in 0.1% sodium citrate for 2 min on ice. Next, TUNEL reaction mixture was added to the cells and incubated at 37°C for 60 min. TMB-positive cells were detected and counted using BD FACSCelesta Flow Cytometer, at least 10,000 cells were analyzed per sample.

### Statistical analysis

Statistical significance was calculated using GraphPad Prism 8.2 package. D’Agostino-Pearson omnibus normality test was used to establish whether or not the population is distributed normally. Unpaired Mann-Whitney test was used to calculate the difference between two different populations which are not normally distributed. One-way analysis of variance (ANOVA) followed by Dunnett’s post hoc test was used for multiple comparisons analysis of normally distributed populations with equal variances (i.e. equal standard deviations (SD)). Brown-Forsythe and Welch ANOVA tests followed by Dunnett’s T3 multiple comparisons analysis were used for normally distributed populations with different SDs. Kruskal-Wallis test followed by followed by Dunn’s multiple comparisons analysis was used for populations which are not normally distributed.

### Data Deposition

The sequence data has been submitted to the Gene Expression Omnibus under superseries identifier GSE152651 which consists of the RNA-Seq data (GSE152650) and the shRNA screen data (GSE152649). Original data and numbers for tables are uploaded to Mendeley Data (DOI: 10.17632/dggchs5s8m.1).

## SUPPLEMENTAL MATERIAL

**Supplemental Figure 1.**
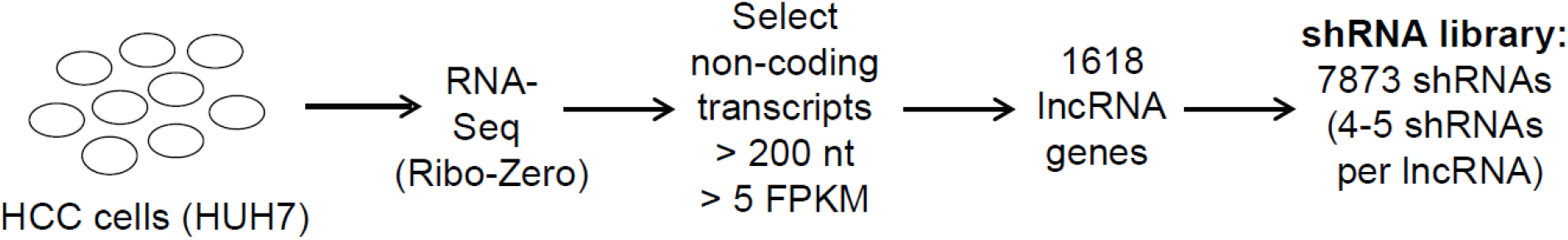
Schematic workflow of shRNA library design.

**Supplemental Figure 2.**
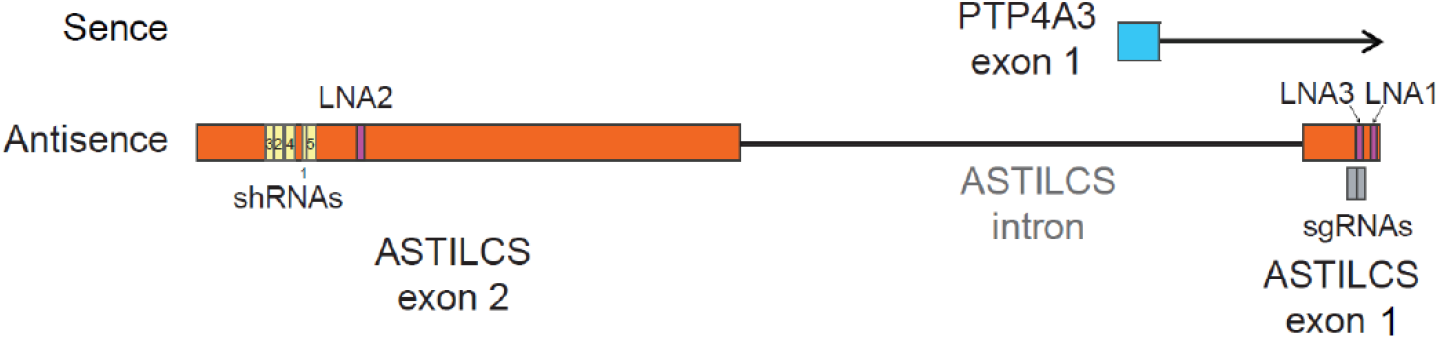
Positions of shRNAs, sgRNAs and LNAs targeting ASTILCS.

**Supplemental Figure 3.**
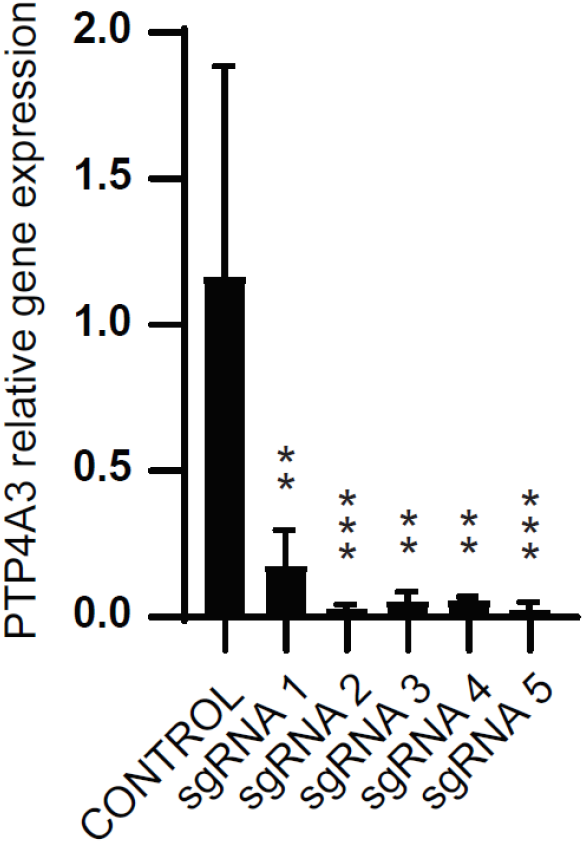
**PTP4A3 expression in HUH7 cells transduced with sgRNAs targeting ASTILCS TSS.** n=12. All values are mean ± SD, **** p < 0.0001

**Supplemental Figure 4.**
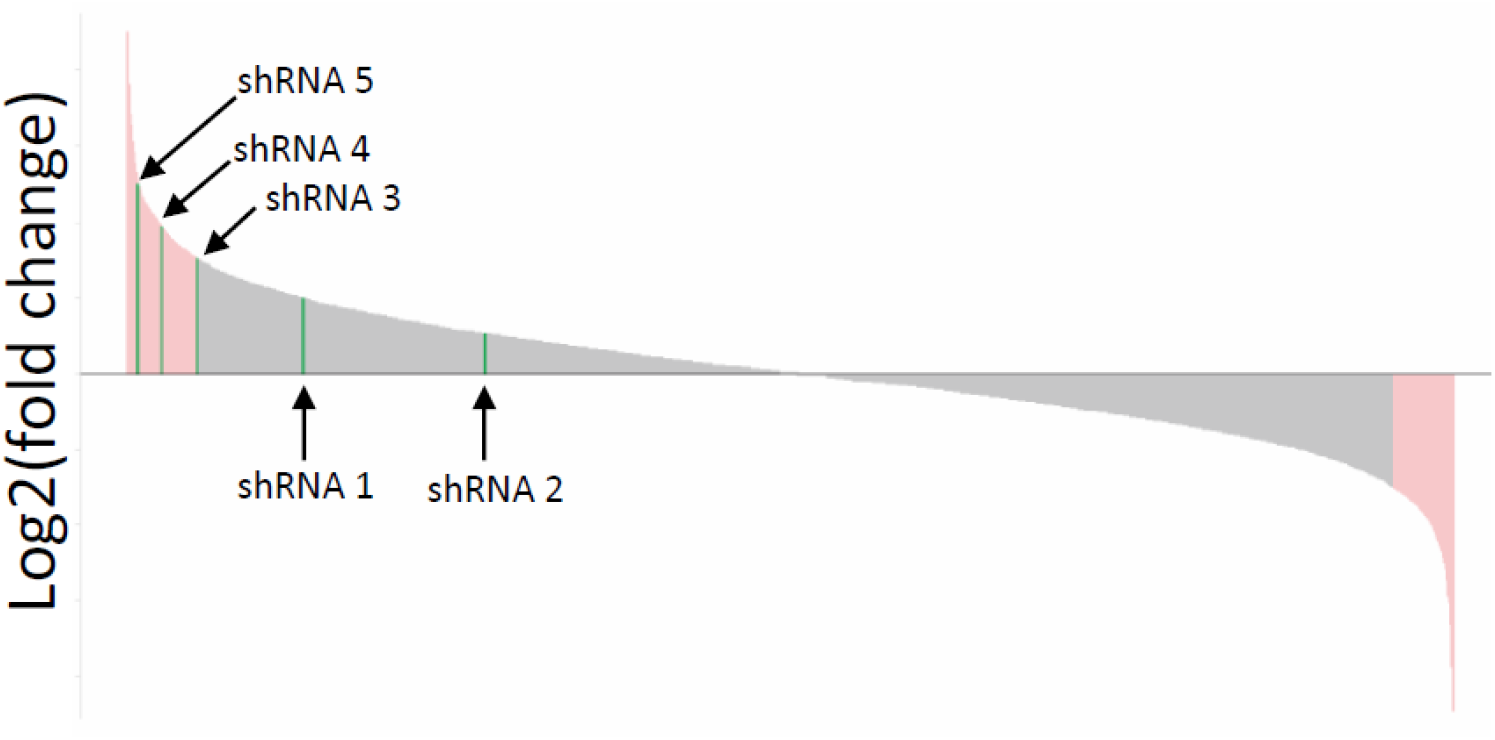
Waterfall plot of shRNAs present in the final population of HUH7 cells. Log2(fold change) > or < 0.75 is highlighted in light pink, shRNAs targeting ASTILCS are highlighted in green.

**Supplemental Figure 5.**
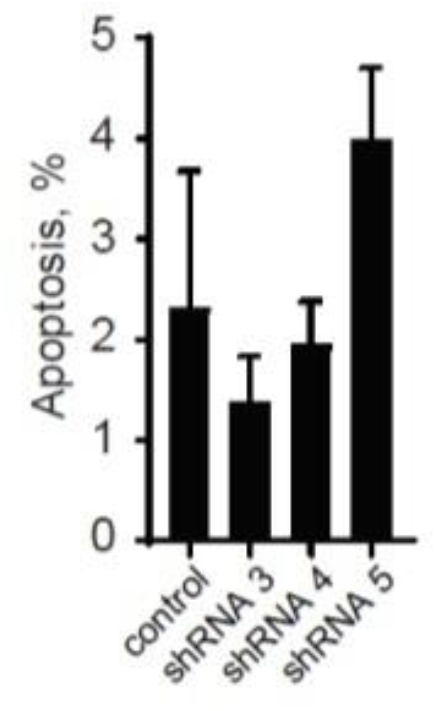
Apoptosis in HUH7 cells treated with shRNAs targeting ASTILCS, n=3. All values are mean ± SD, there is no significant difference compared to control.

**Supplemental Figure 6.**
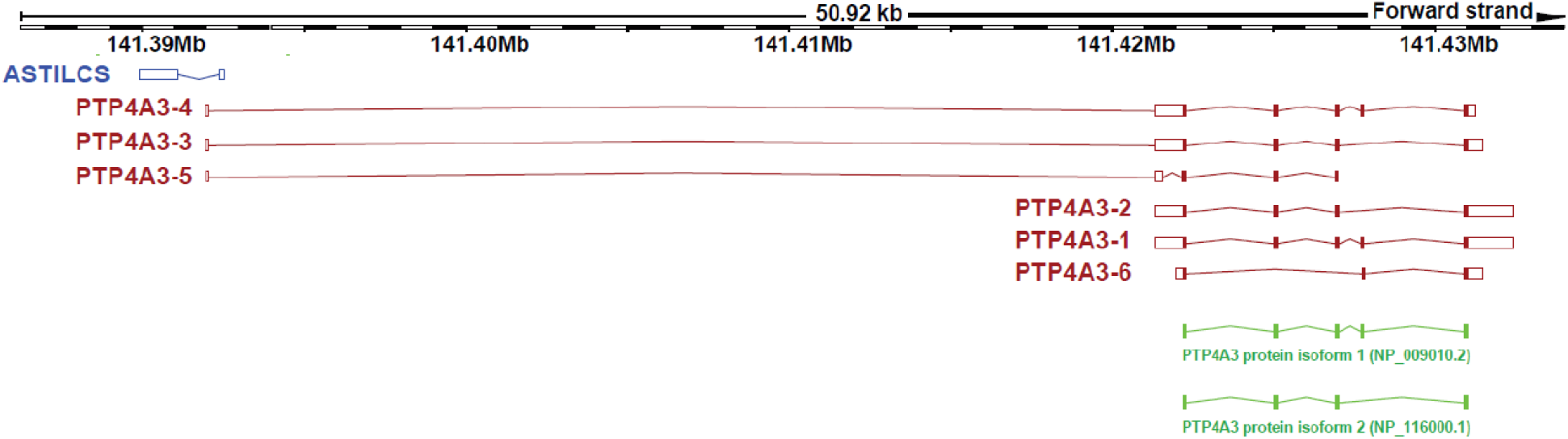
PTP4A3 gene produces 6 transcripts and 2 protein isoforms. Adapted from http://www.ensembl.org/.

**Supplemental Figure 7.**
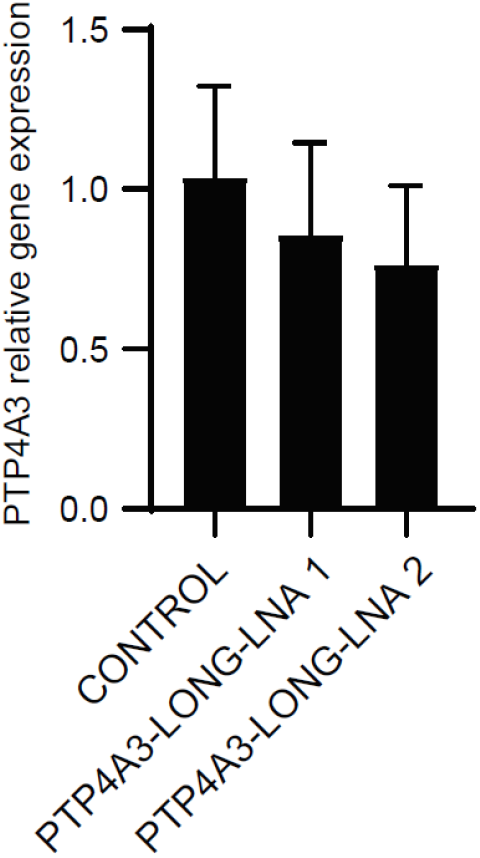
Expression of short PTP4A3 transcripts upon LNA-mediated knockdown of long PTP4A3 transcripts, n≥8. All values are mean ± SD, there is no significant difference compared to control.

**Supplemental Table 1.**
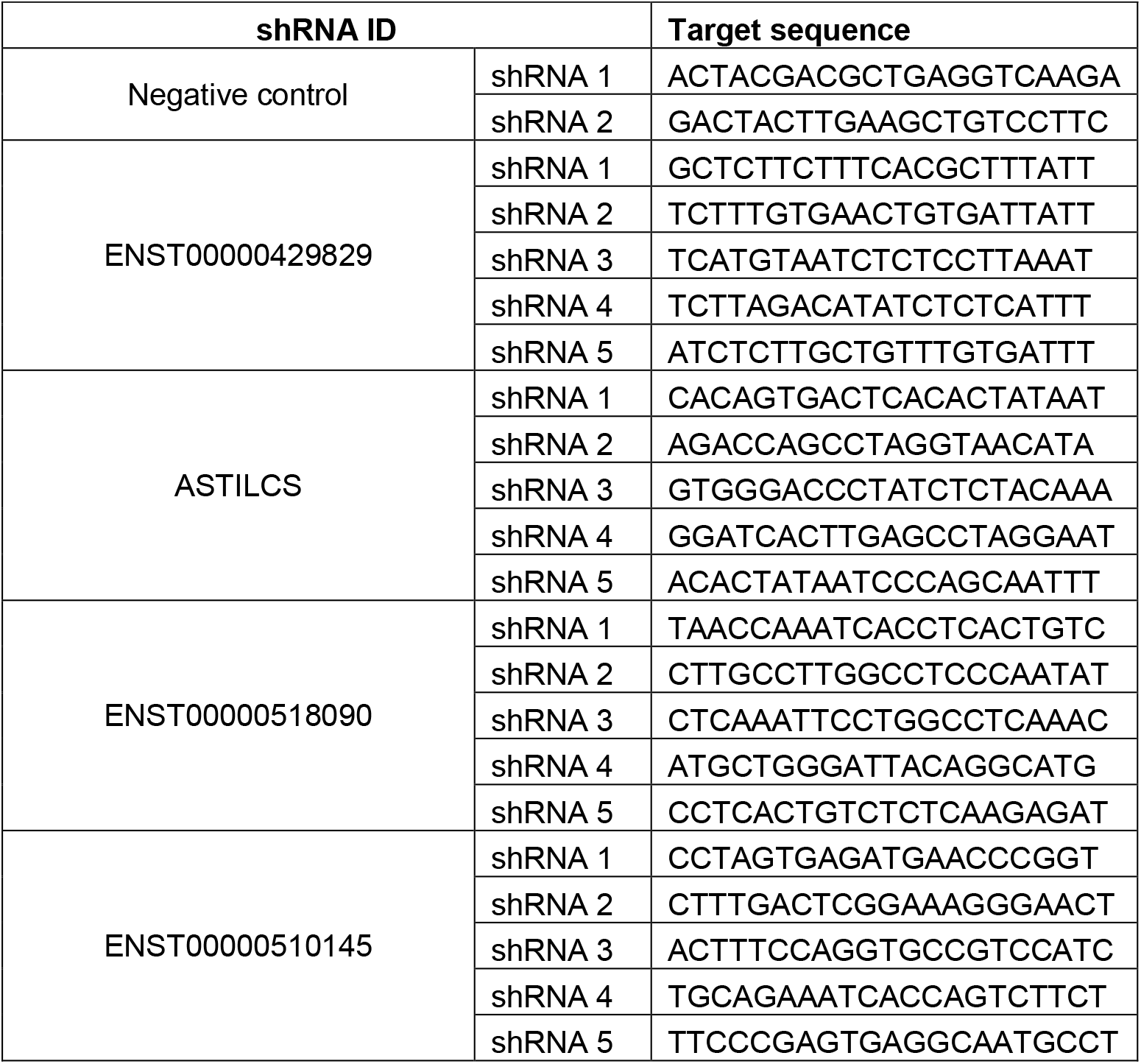
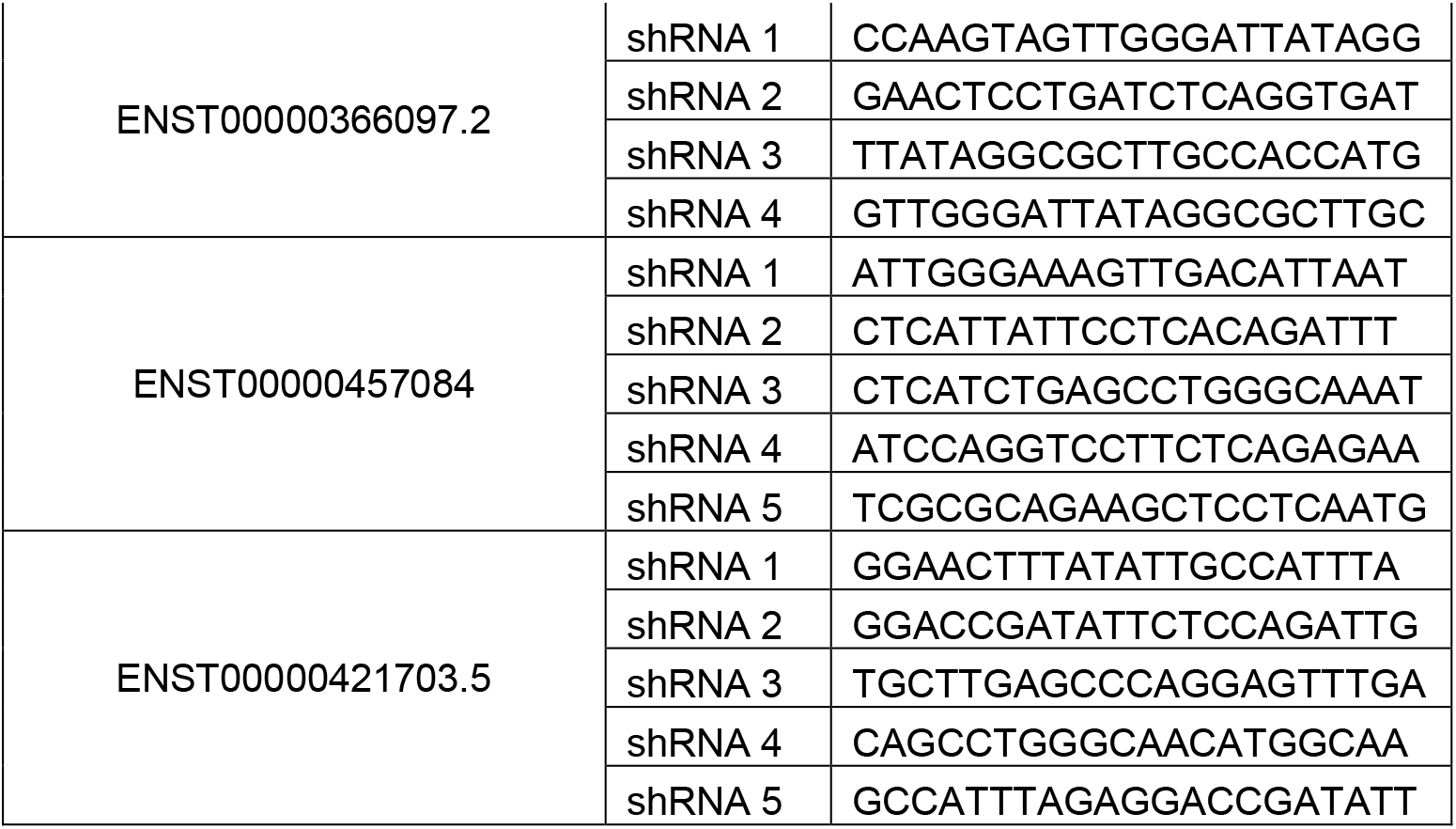
Long non-coding RNA transcripts expressed in HUH7 cells. (Excel file) ***Supplemental Table 2**. shRNAs used for validation of the screen results*

**Supplemental Table 3.**
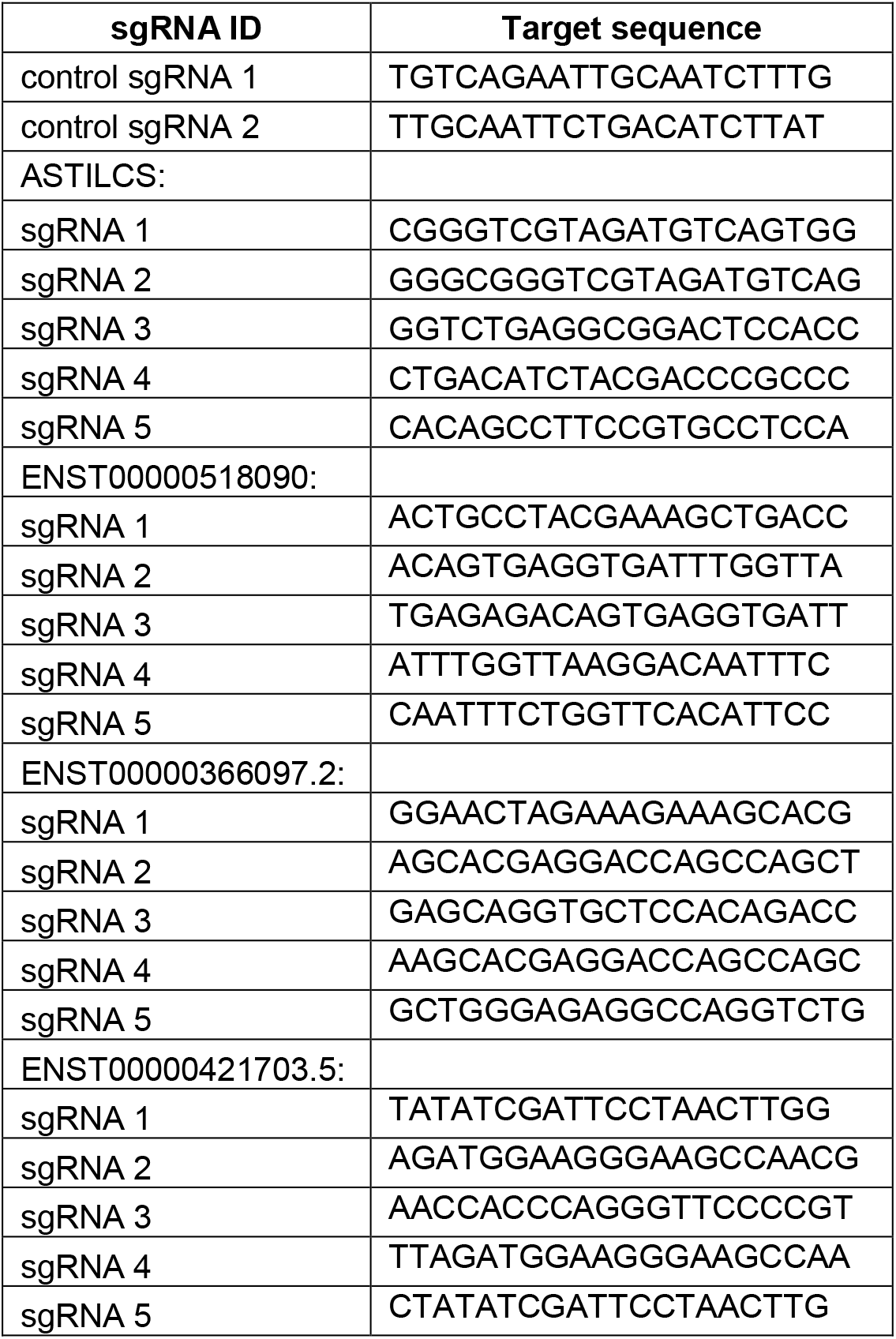
sgRNAs used in the study

**Supplemental Table 4.**
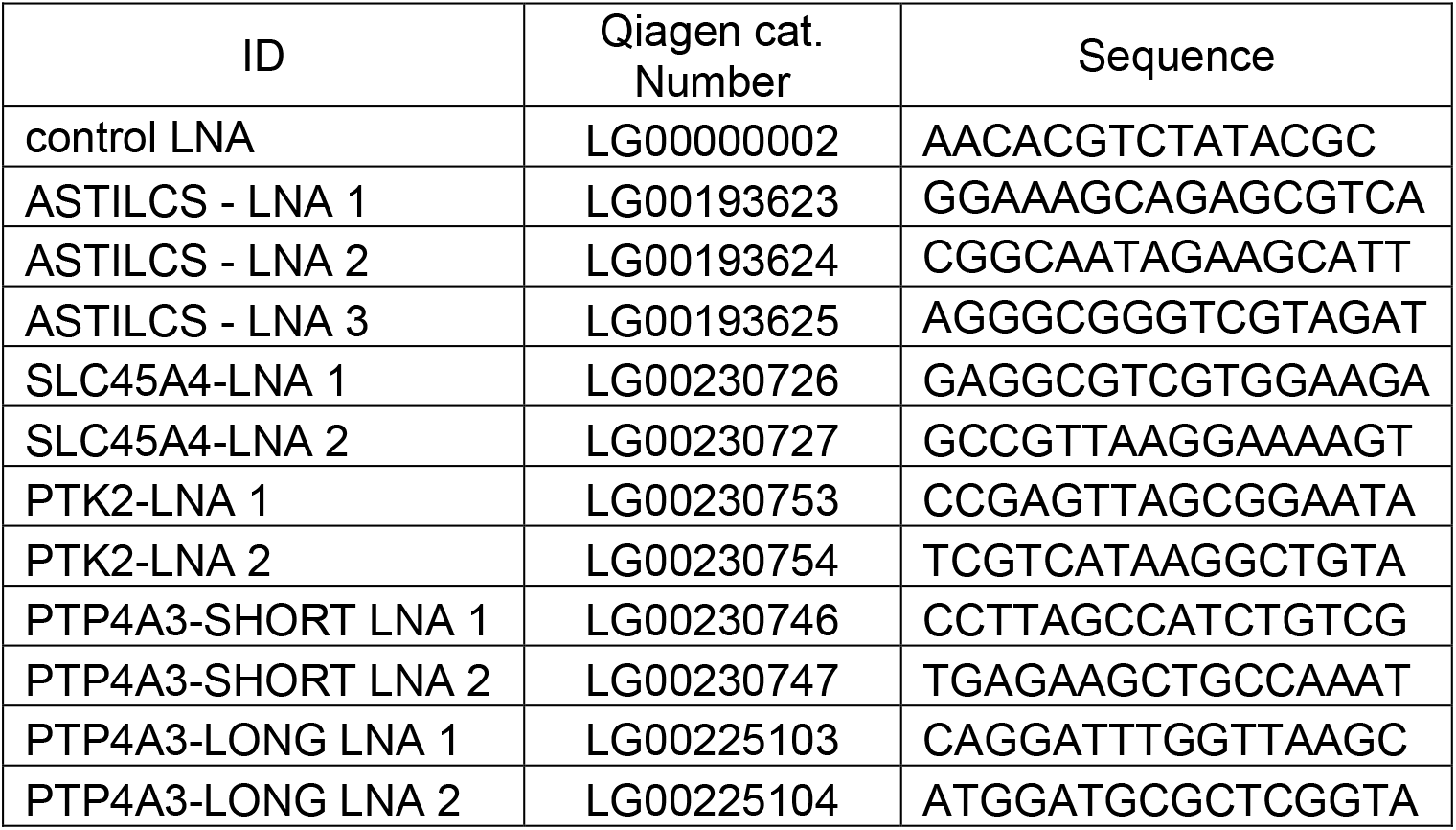
LNAs used in the study

**Supplemental Table 5.** *Population* doubling time (Td) for HUH7 cells expressing GFP and ASTILCS. Td was calculated from the exponential portion of the cell growth curve (days 3-5) using the following equation: Td = 0.693t/ln(Nt/N0), where t–time (in days), N0–initial cell number, Nt–cell number on day t.

|  | control | ASTILCS |
| --- | --- | --- |
| replicate 1 | 1.17 | 1.16 |
| replicate 2 | 1.07 | 1.13 |
| replicate 3 | 1.02 | 1.10 |
| replicate 4 | 1.02 | 1.11 |
| average | 1.07 | 1.13 |
| Stand. Dev. | 0.07 | 0.03 |
|  | p=0.1698 |  |

**Supplemental Table 6.**
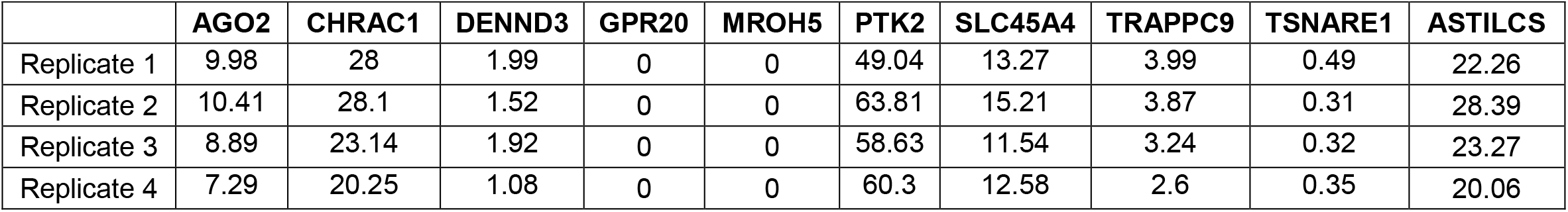
Expression (in FPKM) of genes in the ASTILCS locus in HUH7 cells.

**Supplemental Table 7.**
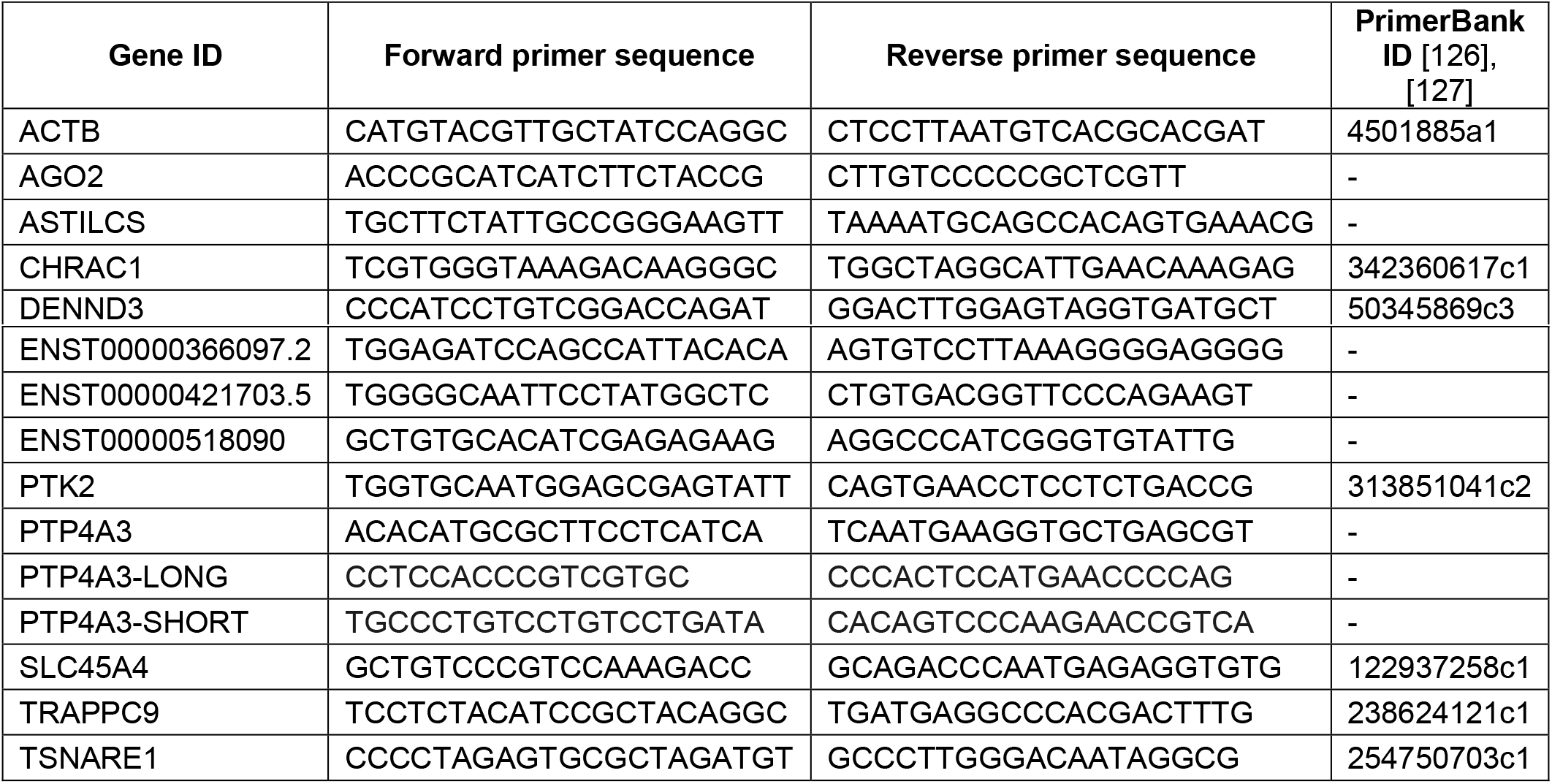
SYBR Green primers used in the study.

## REFERENCES

[1] S. Asia, S. Asia, and H. Hdi, “Liver Cancer Global WHO Report,” vol. 876, pp. 2018—2019, 2018, [Online]. Available: http://gco.iarc.fr/today.

[2] C. Allemani et al., “Global surveillance of cancer survival 1995-2009: Analysis of individual data for 25 676 887 patients from 279 population-based registries in 67 countries (CONCORD-2),” Lancet, vol. 385, no. 9972, pp. 977–1010, 2015, doi: 10.1016/S0140-6736(14)62038-9.

[3] C. Allemani et al., “Global surveillance of trends in cancer survival 2000–14 (CONCORD-3): analysis of individual records for 37 513 025 patients diagnosed with one of 18 cancers from 322 population-based registries in 71 countries,” Lancet, vol. 391, no. 10125, pp. 1023–1075, 2018, doi: 10.1016/S0140-6736(17)33326-3.

[4] A. Villanueva, “Hepatocellular carcinoma,” N. Engl. J. Med., vol. 380, no. 15, pp. 1450–1462, 2019, doi: 10.1056/NEJMra1713263.

[5] P. Rawla, T. Sunkara, P. Muralidharan, and J. P. Raj, “Update in global trends and aetiology of hepatocellular carcinoma,” Współczesna Onkol., vol. 22, no. 3, pp. 141–150, 2018, doi: 10.5114/wo.2018.78941.

[6] American Cancer Society, “Cancer Facts and Figures 2020,” Accessed: 28-Mar-2020. [Online]. Available: https://www.cancer.org/content/dam/cancer-org/research/cancer-facts-and-statistics/annual-cancer-facts-and-figures/2020/cancer-facts-and-figures-2020.pdf.

[7] C. Allain, G. Angenard, B. Clément, and C. Coulouarn, “Integrative Genomic Analysis Identifies the Core Transcriptional Hallmarks of Human Hepatocellular Carcinoma.,” Cancer Res., vol. 76, no. 21, pp. 6374–6381, Nov. 2016, doi: 10.1158/0008-5472.CAN-16-1559.

[8] K. Okrah et al., “Transcriptomic analysis of hepatocellular carcinoma reveals molecular features of disease progression and tumor immune biology,” npj Precis. Oncol., vol. 2, no. 1, 2018, doi: 10.1038/s41698-018-0068-8.

[9] A. Ally et al., “Comprehensive and Integrative Genomic Characterization of Hepatocellular Carcinoma,” Cell, vol. 169, no. 7, pp. 1327–1341.e23, 2017, doi: 10.1016/j.cell.2017.05.046.

[10] A. Fujimoto et al., “Whole-genome sequencing of liver cancers identifies etiological influences on mutation patterns and recurrent mutations in chromatin regulators,” Nat. Genet., vol. 44, no. 7, pp. 760–764, 2012, doi: 10.1038/ng.2291.

[11] Z. Kan et al., “Whole-genome sequencing identifies recurrent mutations in hepatocellular carcinoma,” Genome Res., vol. 23, no. 9, pp. 1422–1433, 2013, doi: 10.1101/gr.154492.113.

[12] K. Schulze et al., “Exome sequencing of hepatocellular carcinomas identifies new mutational signatures and potential therapeutic targets,” Nat. Genet., vol. 47, no. 5, pp. 505–511, 2015, doi: 10.1038/ng.3252.

[13] H. Nakagawa, M. Fujita, and A. Fujimoto, “Genome sequencing analysis of liver cancer for precision medicine,” Semin. Cancer Biol., vol. 55, no. November 2017, pp. 120–127, 2019, doi: 10.1016/j.semcancer.2018.03.004.

[14] K. Chaudhary, O. B. Poirion, L. Lu, S. Huang, T. Ching, and L. X. Garmire, “Multimodal meta-analysis of 1,494 hepatocellular carcinoma samples reveals significant impact of consensus driver genes on phenotypes,” Clin. Cancer Res., vol. 25, no. 2, pp. 463–472, 2019, doi: 10.1158/1078-0432.CCR-18-0088.

[15] J. J. Harding et al., “Prospective genotyping of hepatocellular carcinoma: Clinical implications of next-generation sequencing for matching patients to targeted and immune therapies,” Clin. Cancer Res., vol. 25, no. 7, pp. 2116–2126, 2019, doi: 10.1158/1078-0432.CCR-18-2293.

[16] Y. Hoshida et al., “Integrative transcriptome analysis reveals common molecular subclasses of human hepatocellular carcinoma,” Cancer Res., vol. 69, no. 18, pp. 7385–7392, 2009, doi: 10.1158/0008-5472.CAN-09-1089.

[17] K. Schulze, J. C. Nault, and A. Villanueva, “Genetic profiling of hepatocellular carcinoma using next-generation sequencing,” J. Hepatol., vol. 65, no. 5, pp. 1031–1042, 2016, doi: 10.1016/j.jhep.2016.05.035.

[18] G. Khemlina, S. Ikeda, and R. Kurzrock, “The biology of Hepatocellular carcinoma: Implications for genomic and immune therapies,” Mol. Cancer, vol. 16, no. 1, pp. 1–10, 2017, doi: 10.1186/s12943-017-0712-x.

[19] “Identification and analysis of functional elements in 1% of the human genome by the ENCODE pilot project,” Nature, vol. 447, no. 7146, pp. 799–816, Jun. 2007, doi: 10.1038/nature05874.

[20] “An integrated encyclopedia of DNA elements in the human genome,” Nature, vol. 489, no. 7414, pp. 57–74, Sep. 2012, doi: 10.1038/nature11247.

[21] T. Derrien et al., “The GENCODE v7 catalog of human long noncoding RNAs: Analysis of their gene structure, evolution, and expression,” Genome Res., vol. 22, no. 9, pp. 1775–1789, Sep. 2012, doi: 10.1101/gr.132159.111.

[22] H. Jia, M. Osak, G. K. Bogu, L. W. Stanton, R. Johnson, and L. Lipovich, “Genome-wide computational identification and manual annotation of human long noncoding RNA genes,” RNA, vol. 16, no. 8, pp. 1478–1487, Aug. 2010, doi: 10.1261/rna.1951310.

[23] M. N. Cabili et al., “Integrative annotation of human large intergenic noncoding RNAs reveals global properties and specific subclasses.,” Genes Dev., vol. 25, no. 18, pp. 1915–27, Sep. 2011, doi: 10.1101/gad.17446611.

[24] M. K. Iyer et al., “The landscape of long noncoding RNAs in the human transcriptome,” Nat. Genet., vol. 47, no. 3, pp. 199–208, Mar. 2015, doi: 10.1038/ng.3192.

[25] M. Guttman et al., “Chromatin signature reveals over a thousand highly conserved large non-coding RNAs in mammals,” Nature, vol. 458, no. 7235, pp. 223–227, Mar. 2009, doi: 10.1038/nature07672.

[26] Y. Yang et al., “Recurrently deregulated lncRNAs in hepatocellular carcinoma,” Nat. Commun., vol. 8, 2017, doi: 10.1038/ncomms14421.

[27] Q. Xia et al., “Identification of novel biomarkers for hepatocellular carcinoma using transcriptome analysis,” J. Cell. Physiol., vol. 234, no. 4, pp. 4851–4863, 2019, doi: 10.1002/jcp.27283.

[28] H. Cui, Y. Zhang, Q. Zhang, W. Chen, H. Zhao, and J. Liang, “A comprehensive genome-wide analysis of long noncoding RNA expression profile in hepatocellular carcinoma,” Cancer Med., vol. 6, no. 12, pp. 2932–2941, 2017, doi: 10.1002/cam4.1180.

[29] D. D. Esposti et al., “Identification of novel long non-coding RNAs deregulated in hepatocellular carcinoma using RNA-sequencing,” Oncotarget, vol. 7, no. 22, pp. 31862–31877, 2016, doi: 10.18632/oncotarget.7364.

[30] C.-M. Wong, F. H.-C. Tsang, and I. O.-L. Ng, “Non-coding RNAs in hepatocellular carcinoma: molecular functions and pathological implications.,” Nat. Rev. Gastroenterol. Hepatol., vol. 15, no. 3, pp. 137–151, Mar. 2018, doi: 10.1038/nrgastro.2017.169.

[31] X. Huo et al., “Dysregulated long noncoding RNAs (lncRNAs) in hepatocellular carcinoma: implications for tumorigenesis, disease progression, and liver cancer stem cells.,” Mol. Cancer, vol. 16, no. 1, p. 165, Dec. 2017, doi: 10.1186/s12943-017-0734-4.

[32] M. Klingenberg, A. Matsuda, S. Diederichs, and T. Patel, “Non-coding RNA in hepatocellular carcinoma: Mechanisms, biomarkers and therapeutic targets,” J. Hepatol., vol. 67, no. 3, pp. 603–618, Sep. 2017, doi: 10.1016/j.jhep.2017.04.009.

[33] P. McDonel and M. Guttman, “Approaches for Understanding the Mechanisms of Long Noncoding RNA Regulation of Gene Expression.,” Cold Spring Harb. Perspect. Biol., vol. 11, no. 12, p. a032151, Dec. 2019, doi: 10.1101/cshperspect.a032151.

[34] L.-L. Chen, “Linking Long Noncoding RNA Localization and Function,” Trends Biochem. Sci., vol. 41, no. 9, pp. 761–772, Sep. 2016, doi: 10.1016/j.tibs.2016.07.003.

[35] J. Carlevaro-Fita and R. Johnson, “Global Positioning System: Understanding Long Noncoding RNAs through Subcellular Localization,” Mol. Cell, vol. 73, no. 5, pp. 869–883, Mar. 2019, doi: 10.1016/j.molcel.2019.02.008.

[36] J. M. Engreitz et al., “Local regulation of gene expression by lncRNA promoters, transcription and splicing,” Nature, vol. 539, no. 7629, pp. 452–455, Nov. 2016, doi: 10.1038/nature20149.

[37] J. H. Noh, K. M. Kim, W. G. McClusky, K. Abdelmohsen, and M. Gorospe, “Cytoplasmic functions of long noncoding RNAs,” Wiley Interdiscip. Rev. RNA, vol. 9, no. 3, p. e1471, May 2018, doi: 10.1002/wrna.1471.

[38] D. E. Root, N. Hacohen, W. C. Hahn, E. S. Lander, and D. M. Sabatini, “Genome-scale loss-of-function screening with a lentiviral RNAi library,” Nat. Methods, vol. 3, no. 9, pp. 715–719, Sep. 2006, doi: 10.1038/nmeth924.

[39] L. A. Gilbert et al., “Genome-Scale CRISPR-Mediated Control of Gene Repression and Activation,” Cell, vol. 159, no. 3, pp. 647–661, Oct. 2014, doi: 10.1016/j.cell.2014.09.029.

[40] M. Kampmann et al., “Next-generation libraries for robust RNA interference-based genome-wide screens.,” Proc. Natl. Acad. Sci., vol. 112, no. 26, pp. E3384–91, Jun. 2015, doi: 10.1073/pnas.1508821112.

[41] J. Beermann et al., “A large shRNA library approach identifies lncRNA Ntep as an essential regulator of cell proliferation,” Cell Death Differ., vol. 25, no. 2, pp. 307–318, Feb. 2018, doi: 10.1038/cdd.2017.158.

[42] J.-F. Huang et al., “Genome-wide screening identifies oncofetal lncRNA Ptn-dt promoting the proliferation of hepatocellular carcinoma cells by regulating the Ptn receptor,” Oncogene, vol. 38, no. 18, pp. 3428–3445, May 2019, doi: 10.1038/s41388-018-0643-z.

[43] J. Joung et al., “Genome-scale activation screen identifies a lncRNA locus regulating a gene neighbourhood,” Nature, vol. 548, no. 7667, pp. 343–346, Aug. 2017, doi: 10.1038/nature23451.

[44] I. Tiessen et al., “A high-throughput screen identifies the long non-coding RNA DRAIC as a regulator of autophagy,” Oncogene, vol. 38, no. 26, pp. 5127–5141, Jun. 2019, doi: 10.1038/s41388-019-0783-9.

[45] P. Cai et al., “A genome-wide long noncoding RNA CRISPRi screen identifies *PRANCR* as a novel regulator of epidermal homeostasis,” Genome Res., vol. 30, no. 1, pp. 22–34, Jan. 2020, doi: 10.1101/gr.251561.119.

[46] R. Galeev et al., “Genome-wide RNAi Screen Identifies Cohesin Genes as Modifiers of Renewal and Differentiation in Human HSCs,” Cell Rep., vol. 14, no. 12, pp. 2988–3000, Mar. 2016, doi: 10.1016/j.celrep.2016.02.082.

[47] S. E. Castel and R. A. Martienssen, “RNA interference in the nucleus: roles for small RNAs in transcription, epigenetics and beyond,” Nat. Rev. Genet., vol. 14, no. 2, pp. 100–112, Feb. 2013, doi: 10.1038/nrg3355.

[48] R. Kalantari, C.-M. Chiang, and D. R. Corey, “Regulation of mammalian transcription and splicing by Nuclear RNAi,” Nucleic Acids Res., vol. 44, no. 2, pp. 524–537, Jan. 2016, doi: 10.1093/nar/gkv1305.

[49] K. A. Lennox and M. A. Behlke, “Cellular localization of long non-coding RNAs affects silencing by RNAi more than by antisense oligonucleotides,” Nucleic Acids Res., vol. 44, no. 2, pp. 863–877, Jan. 2016, doi: 10.1093/nar/gkv1206.

[50] S. Avivi et al., “Visualizing nuclear RNAi activity in single living human cells,” Proc. Natl. Acad. Sci. U. S. A., vol. 114, no. 42, pp. E8837–E8846, 2017, doi: 10.1073/pnas.1707440114.

[51] L. S. Qi et al., “Repurposing CRISPR as an RNA-Guided Platform for Sequence-Specific Control of Gene Expression,” Cell, vol. 152, no. 5, pp. 1173–1183, Feb. 2013, doi: 10.1016/j.cell.2013.02.022.

[52] P. I. Thakore et al., “Highly specific epigenome editing by CRISPR-Cas9 repressors for silencing of distal regulatory elements,” Nat. Methods, vol. 12, no. 12, pp. 1143–1149, Dec. 2015, doi: 10.1038/nmeth.3630.

[53] A. Goyal, K. Myacheva, M. Groß, M. Klingenberg, B. Duran Arqué, and S. Diederichs, “Challenges of CRISPR/Cas9 applications for long non-coding RNA genes,” Nucleic Acids Res., p. gkw883, Sep. 2016, doi: 10.1093/nar/gkw883.

[54] X. Q. Lin, Z. M. Huang, X. Chen, F. Wu, and W. Wu, “XIST Induced by JPX Suppresses Hepatocellular Carcinoma by Sponging miR-155-5p.,” Yonsei Med. J., vol. 59, no. 7, pp. 816–826, Sep. 2018, doi: 10.3349/ymj.2018.59.7.816.

[55] Q. Kong et al., “LncRNA XIST functions as a molecular sponge of miR-194-5p to regulate MAPK1 expression in hepatocellular carcinoma cell,” J. Cell. Biochem., vol. 119, no. 6, pp. 4458–4468, Jun. 2018, doi: 10.1002/jcb.26540.

[56] S. Chang, B. Chen, X. Wang, K. Wu, and Y. Sun, “Long non-coding RNA XIST regulates PTEN expression by sponging miR-181a and promotes hepatocellular carcinoma progression.,” BMC Cancer, vol. 17, no. 1, p. 248, Dec. 2017, doi: 10.1186/s12885-017-3216-6.

[57] W. Ma et al., “Downregulation of long non-coding RNAs JPX and XIST is associated with the prognosis of hepatocellular carcinoma.,” Clin. Res. Hepatol. Gastroenterol., vol. 41, no. 2, pp. 163–170, Mar. 2017, doi: 10.1016/j.clinre.2016.09.002.

[58] L. K. Zhuang et al., “MicroRNA-92b promotes hepatocellular carcinoma progression by targeting Smad7 and is mediated by long non-coding RNA XIST,” Cell Death Dis., vol. 7, no. 4, pp. e2203–e2203, Apr. 2016, doi: 10.1038/cddis.2016.100.

[59] J. Wan, D. Deng, X. Wang, X. Wang, S. Jiang, and R. Cui, “LINC00491 as a new molecular marker can promote the proliferation, migration and invasion of colon adenocarcinoma cells,” Onco. Targets. Ther., vol. 12, pp. 6471–6480, 2019, doi: 10.2147/OTT.S201233.

[60] J. Li et al., “Long non-coding RNAs expressed in pancreatic ductal adenocarcinoma and lncRNA BC008363 an independent prognostic factor in PDAC,” Pancreatology, vol. 14, no. 5, pp. 385–390, Sep. 2014, doi: 10.1016/J.PAN.2014.07.013.

[61] A. Sahakyan, Y. Yang, and K. Plath, “The Role of Xist in X-Chromosome Dosage Compensation,” Trends Cell Biol., vol. 28, no. 12, pp. 999–1013, Dec. 2018, doi: 10.1016/j.tcb.2018.05.005.

[62] D. Chen et al., “Long noncoding RNA XIST expedites metastasis and modulates epithelial–mesenchymal transition in colorectal cancer,” Cell Death Dis., vol. 8, no. 8, pp. e3011–e3011, Aug. 2017, doi: 10.1038/cddis.2017.421.

[63] C. Li et al., “Long non-coding RNA XIST promotes TGF-β-induced epithelial-mesenchymal transition by regulating miR-367/141-ZEB2 axis in non-small-cell lung cancer,” Cancer Lett., vol. 418, pp. 185–195, Apr. 2018, doi: 10.1016/j.canlet.2018.01.036.

[64] F. Xing et al., “Loss of XIST in Breast Cancer Activates MSN-c-Met and Reprograms Microglia via Exosomal miRNA to Promote Brain Metastasis,” Cancer Res., vol. 78, no. 15, pp. 4316–4330, Aug. 2018, doi: 10.1158/0008-5472.CAN-18-1102.

[65] C. J. Echeverri and N. Perrimon, “High-throughput RNAi screening in cultured cells: a user’s guide,” Nat. Rev. Genet., vol. 7, no. 5, pp. 373–384, May 2006, doi: 10.1038/nrg1836.

[66] W. G. Kaelin and Jr, “Use and Abuse of RNAi to Study Mammalian Gene Function: Molecular Biology,” Science, vol. 337, no. 6093, p. 421, 2012, doi: 10.1126/SCIENCE.1225787.

[67] P.-J. Volders et al., “LNCipedia 5: towards a reference set of human long non-coding RNAs,” Nucleic Acids Res., vol. 47, no. D1, pp. D135–D139, Jan. 2019, doi: 10.1093/nar/gky1031.

[68] J. Li et al., “TANRIC: An Interactive Open Platform to Explore the Function of lncRNAs in Cancer,” Cancer Res., vol. 75, no. 18, pp. 3728–3737, Sep. 2015, doi: 10.1158/0008-5472.CAN-15-0273.

[69] T. S. Slordahl et al., “The phosphatase of regenerating liver-3 (PRL-3) is important for IL-6-mediated survival of myeloma cells.,” Oncotarget, vol. 7, no. 19, pp. 27295–27306, May 2016, doi: 10.18632/oncotarget.8422.

[70] Y.-X. Lian, R. Chen, Y.-H. Xu, C.-L. Peng, and H.-C. Hu, “Effect of protein-tyrosine phosphatase 4A3 by small interfering RNA on the proliferation of lung cancer.,” Gene, vol. 511, no. 2, pp. 169–176, Dec. 2012, doi: 10.1016/j.gene.2012.09.079.

[71] J. Zhang et al., “miR-21, miR-17 and miR-19a induced by phosphatase of regenerating liver-3 promote the proliferation and metastasis of colon cancer.,” Br. J. Cancer, vol. 107, no. 2, pp. 352–359, Jul. 2012, doi: 10.1038/bjc.2012.251.

[72] Y. Matsukawa, S. Semba, H. Kato, Y.-I. Koma, K. Yanagihara, and H. Yokozaki, “Constitutive suppression of PRL-3 inhibits invasion and proliferation of gastric cancer cell in vitro and in vivo.,” Pathobiology, vol. 77, no. 3, pp. 155–162, 2010, doi: 10.1159/000292649.

[73] L. Wang et al., “PTP4A3 is a target for inhibition of cell proliferatin, migration and invasion through Akt/mTOR signaling pathway in glioblastoma under the regulation of miR-137.,” Brain Res., vol. 1646, pp. 441–450, Sep. 2016, doi: 10.1016/j.brainres.2016.06.026.

[74] J. Minshull and T. Hunt, “The use of single-stranded DNA and RNase H to promote quantitative ‘hybrid arrest of translation’ of mRNA/DNA hybrids in reticulocyte lysate cell-free translations,” Nucleic Acids Res., vol. 14, no. 16, pp. 6433–6451, 1986, doi: 10.1093/nar/14.16.6433.

[75] H. Nakamura et al., “How does RNase H recognize a DNA.RNA hybrid?,” Proc. Natl. Acad. Sci., vol. 88, no. 24, pp. 11535–11539, Dec. 1991, doi: 10.1073/pnas.88.24.11535.

[76] T. A. Vickers and S. T. Crooke, “The rates of the major steps in the molecular mechanism of RNase H1-dependent antisense oligonucleotide induced degradation of RNA,” Nucleic Acids Res., vol. 43, no. 18, pp. 8955–8963, Oct. 2015, doi: 10.1093/nar/gkv920.

[77] G.-P. C. Mologni L, leCoutre P, Nielsen PE, “Additive antisense effects of different PNAs on the in vitro translation of the PML/RARalpha gene,” Nucleic Acids Res., vol. 26, no. 8, pp. 1934–1938, Apr. 1998, doi: 10.1093/nar/26.8.1934.

[78] B. F. Baker et al., “2’-*O* -(2-Methoxy)ethyl-modified Anti-intercellular Adhesion Molecule 1 (ICAM-1) Oligonucleotides Selectively Increase the ICAM-1 mRNA Level and Inhibit Formation of the ICAM-1 Translation Initiation Complex in Human Umbilical Vein Endothelial Cell,” J. Biol. Chem., vol. 272, no. 18, pp. 11994–12000, May 1997, doi: 10.1074/jbc.272.18.11994.

[79] N. Dias and C. A. Stein, “Antisense Oligonucleotides: Basic Concepts and Mechanisms,” Mol. Cancer Ther., vol. 1, no. 5, pp. 347–355, Mar. 2002, Accessed: 03-May-2020. [Online]. Available: https://mct.aacrjournals.org/content/1/5/347.long.

[80] C. F. Bennett and E. E. Swayze, “RNA Targeting Therapeutics: Molecular Mechanisms of Antisense Oligonucleotides as a Therapeutic Platform,” Annu. Rev. Pharmacol. Toxicol., vol. 50, no. 1, pp. 259–293, Feb. 2010, doi: 10.1146/annurev.pharmtox.010909.105654.

[81] R. Kole, A. R. Krainer, and S. Altman, “RNA therapeutics: beyond RNA interference and antisense oligonucleotides,” Nat. Rev. Drug Discov., vol. 11, no. 2, pp. 125–140, Feb. 2012, doi: 10.1038/nrd3625.

[82] N. Gil and I. Ulitsky, “Regulation of gene expression by cis-acting long non-coding RNAs,” Nat. Rev. Genet., vol. 21, no. 2, pp. 102–117, Feb. 2020, doi: 10.1038/s41576-019-0184-5.

[83] F. Kopp and J. T. Mendell, “Functional Classification and Experimental Dissection of Long Noncoding RNAs,” Cell, vol. 172, no. 3, pp. 393–407, Jan. 2018, doi: 10.1016/j.cell.2018.01.011.

[84] O. Khorkova, A. J. Myers, J. Hsiao, and C. Wahlestedt, “Natural antisense transcripts,” Hum. Mol. Genet., vol. 23, no. R1, pp. R54–R63, Sep. 2014, doi: 10.1093/hmg/ddu207.

[85] V. Pelechano and L. M. Steinmetz, “Gene regulation by antisense transcription,” Nat. Rev. Genet., vol. 14, no. 12, pp. 880–893, Dec. 2013, doi: 10.1038/nrg3594.

[86] D. Gnani et al., “Focal adhesion kinase depletion reduces human hepatocellular carcinoma growth by repressing enhancer of zeste homolog 2.,” Cell Death Differ., vol. 24, no. 5, pp. 889–902, May 2017, doi: 10.1038/cdd.2017.34.

[87] M. Lanzafame, G. Bianco, L. M. Terracciano, C. K. Y. Ng, and S. Piscuoglio, “The Role of Long Non-Coding RNAs in Hepatocarcinogenesis.,” Int. J. Mol. Sci., vol. 19, no. 3, p. 682, Feb. 2018, doi: 10.3390/ijms19030682.

[88] C. Li, J. Chen, K. Zhang, B. Feng, R. Wang, and L. Chen, “Progress and Prospects of Long Noncoding RNAs (lncRNAs) in Hepatocellular Carcinoma.,” Cell. Physiol. Biochem., vol. 36, no. 2, pp. 423–434, 2015, doi: 10.1159/000430109.

[89] J. T. Parsons, “Focal adhesion kinase: the first ten years,” J. Cell Sci., vol. 116, no. 8, pp. 1409–1416, Apr. 2003, doi: 10.1242/jcs.00373.

[90] S. K. Mitra, D. A. Hanson, and D. D. Schlaepfer, “Focal adhesion kinase: in command and control of cell motility.,” Nat. Rev. Mol. Cell Biol., vol. 6, no. 1, pp. 56–68, Jan. 2005, doi: 10.1038/nrm1549.

[91] M. D. Schaller, “Cellular functions of FAK kinases: insight into molecular mechanisms and novel functions,” J. Cell Sci., vol. 123, no. 7, pp. 1007–1013, Apr. 2010, doi: 10.1242/jcs.045112.

[92] E. G. Kleinschmidt and D. D. Schlaepfer, “Focal adhesion kinase signaling in unexpected places.,” Curr. Opin. Cell Biol., vol. 45, pp. 24–30, Apr. 2017, doi: 10.1016/j.ceb.2017.01.003.

[93] S.-T. Lim et al., “Nuclear FAK Promotes Cell Proliferation and Survival through FERM-Enhanced p53 Degradation,” Mol. Cell, vol. 29, no. 1, pp. 9–22, Jan. 2008, doi: 10.1016/j.molcel.2007.11.031.

[94] F. J. Sulzmaier, C. Jean, and D. D. Schlaepfer, “FAK in cancer: mechanistic findings and clinical applications,” Nat. Rev. Cancer, vol. 14, no. 9, pp. 598–610, Sep. 2014, doi: 10.1038/nrc3792.

[95] N. Panera, A. Crudele, I. Romito, D. Gnani, and A. Alisi, “Focal Adhesion Kinase: Insight into Molecular Roles and Functions in Hepatocellular Carcinoma.,” Int. J. Mol. Sci., vol. 18, no. 1, p. 99, Jan. 2017, doi: 10.3390/ijms18010099.

[96] T. Shimizu et al., “A first-in-Asian phase 1 study to evaluate safety, pharmacokinetics and clinical activity of VS-6063, a focal adhesion kinase (FAK) inhibitor in Japanese patients with advanced solid tumors.,” Cancer Chemother. Pharmacol., vol. 77, no. 5, pp. 997–1003, May 2016, doi: 10.1007/s00280-016-3010-1.

[97] N. F. Brown et al., “A study of the focal adhesion kinase inhibitor GSK2256098 in patients with recurrent glioblastoma with evaluation of tumor penetration of [11C]GSK2256098,” Neuro. Oncol., vol. 20, no. 12, pp. 1634–1642, Nov. 2018, doi: 10.1093/neuonc/noy078.

[98] S. F. Jones et al., “A phase I study of VS-6063, a second-generation focal adhesion kinase inhibitor, in patients with advanced solid tumors.,” Invest. New Drugs, vol. 33, no. 5, pp. 1100–7, Oct. 2015, doi: 10.1007/s10637-015-0282-y.

[99] Z. Fan et al., “PTK2 promotes cancer stem cell traits in hepatocellular carcinoma by activating Wnt/β-catenin signaling,” Cancer Lett., vol. 450, pp. 132–143, May 2019, doi: 10.1016/j.canlet.2019.02.040.

[100] N. Shang et al., “Focal Adhesion Kinase and β-Catenin Cooperate to Induce Hepatocellular Carcinoma,” Hepatology, vol. 70, no. 5, pp. 1631–1645, Nov. 2019, doi: 10.1002/hep.30707.

[101] T. Fujii et al., “Focal adhesion kinase is overexpressed in hepatocellular carcinoma and can be served as an independent prognostic factor.,” J. Hepatol., vol. 41, no. 1, pp. 104–11, Jul. 2004, doi: 10.1016/j.jhep.2004.03.029.

[102] S. Itoh et al., “Role of expression of focal adhesion kinase in progression of hepatocellular carcinoma.,” Clin. Cancer Res., vol. 10, no. 8, pp. 2812–7, Apr. 2004, doi: 10.1158/1078-0432.ccr-1046-03.

[103] Z. Yuan, Q. Zheng, J. Fan, K. Ai, J. Chen, and X. Huang, “Expression and prognostic significance of focal adhesion kinase in hepatocellular carcinoma,” J. Cancer Res. Clin. Oncol., vol. 136, no. 10, pp. 1489–1496, Oct. 2010, doi: 10.1007/s00432-010-0806-y.

[104] Y.-J. Jan et al., “Overexpressed focal adhesion kinase predicts a higher incidence of extrahepatic metastasis and worse survival in hepatocellular carcinoma.,” Hum. Pathol., vol. 40, no. 10, pp. 1384–90, Oct. 2009, doi: 10.1016/j.humpath.2009.03.006.

[105] H. Okamoto, K. Yasui, C. Zhao, S. Arii, and J. Inazawa, “PTK2 and EIF3S3 genes may be amplification targets at 8q23-q24 and are associated with large hepatocellular carcinomas.,” Hepatology, vol. 38, no. 5, pp. 1242–9, Nov. 2003, doi: 10.1053/jhep.2003.50457.

[106] K. Hashimoto et al., “Analysis of DNA copy number aberrations in hepatitis C virus-associated hepatocellular carcinomas by conventional CGH and array CGH,” Mod. Pathol., vol. 17, no. 6, pp. 617–622, Jun. 2004, doi: 10.1038/modpathol.3800107.

[107] J. Gao et al., “Integrative analysis of complex cancer genomics and clinical profiles using the cBioPortal.,” Sci. Signal., vol. 6, no. 269, p. pl1, Apr. 2013, doi: 10.1126/scisignal.2004088.

[108] Y. Zhao, H. Sun, and H. Wang, “Long noncoding RNAs in DNA methylation: new players stepping into the old game.,” Cell Biosci., vol. 6, no. 1, p. 45, Dec. 2016, doi: 10.1186/s13578-016-0109-3.

[109] J. Zhou, S. Wang, J. Lu, J. Li, and Y. Ding, “Over-expression of phosphatase of regenerating liver-3 correlates with tumor progression and poor prognosis in nasopharyngeal carcinoma.,” Int. J. cancer, vol. 124, no. 8, pp. 1879–86, Apr. 2009, doi: 10.1002/ijc.24096.

[110] W.-B. Zhao, Y. Li, X. Liu, L.-Y. Zhang, and X. Wang, “Evaluation of PRL-3 expression, and its correlation with angiogenesis and invasion in hepatocellular carcinoma.,” Int. J. Mol. Med., vol. 22, no. 2, pp. 187–92, Aug. 2008, Accessed: 06-May-2020. [Online]. Available: http://www.ncbi.nlm.nih.gov/pubmed/18636172.

[111] C. Laurent et al., “High PTP4A3 phosphatase expression correlates with metastatic risk in uveal melanoma patients.,” Cancer Res., vol. 71, no. 3, pp. 666–74, Feb. 2011, doi: 10.1158/0008-5472.CAN-10-0605.

[112] B.-H. Li, Y. Wang, C.-Y. Wang, M.-J. Zhao, T. Deng, and X.-Q. Ren, “Up-Regulation of Phosphatase in Regenerating Liver-3 (PRL-3) Contributes to Malignant Progression of Hepatocellular Carcinoma by Activating Phosphatase and Tensin Homolog Deleted on Chromosome Ten (PTEN)/Phosphoinositide 3-Kinase (PI3K)/AKT Signaling Path,” Med. Sci. Monit., vol. 24, pp. 8105–8114, Nov. 2018, doi: 10.12659/MSM.913307.

[113] B. Langmead, C. Trapnell, M. Pop, and S. L. Salzberg, “Ultrafast and memory-efficient alignment of short DNA sequences to the human genome.,” Genome Biol., vol. 10, no. 3, p. R25, 2009, doi: 10.1186/gb-2009-10-3-r25.

[114] L. B and D. CN, “RSEM: Accurate Transcript Quantification From RNA-Seq Data With or Without a Reference Genome,” BMC Bioinformatics, vol. 12, p. 323, Aug. 2011, doi: 10.1186/1471-2105-12-323.

[115] J. Moffat et al., “A Lentiviral RNAi Library for Human and Mouse Genes Applied to an Arrayed Viral High-Content Screen,” Cell, vol. 124, no. 6, pp. 1283–1298, Mar. 2006, doi: 10.1016/j.cell.2006.01.040.

[116] M. Marcel, Embnet.news: European Molecular Biology Network newsletter., vol. 17, no. 1. EMBnet, Administration Office, 1994.

[117] H. Li and R. Durbin, “Fast and accurate short read alignment with Burrows-Wheeler transform.,” Bioinformatics, vol. 25, no. 14, pp. 1754–60, Jul. 2009, doi: 10.1093/bioinformatics/btp324.

[118] H. Li et al., “The Sequence Alignment/Map format and SAMtools.,” Bioinformatics, vol. 25, no. 16, pp. 2078–9, Aug. 2009, doi: 10.1093/bioinformatics/btp352.

[119] M. I. Love, W. Huber, and S. Anders, “Moderated estimation of fold change and dispersion for RNA-seq data with DESeq2.,” Genome Biol., vol. 15, no. 12, p. 550, 2014, doi: 10.1186/s13059-014-0550-8.

[120] D. Wiederschain et al., “Single-vector inducible lentiviral RNAi system for oncology target validation,” Cell Cycle, vol. 8, no. 3, pp. 498–504, Feb. 2009, doi: 10.4161/cc.8.3.7701.

[121] J. G. Doench et al., “Optimized sgRNA design to maximize activity and minimize off-target effects of CRISPR-Cas9,” Nat. Biotechnol., vol. 34, no. 2, pp. 184–191, Feb. 2016, doi: 10.1038/nbt.3437.

[122] K. R. Sanson et al., “Optimized libraries for CRISPR-Cas9 genetic screens with multiple modalities,” Nat. Commun., vol. 9, no. 1, p. 5416, Dec. 2018, doi: 10.1038/s41467-018-07901-8.

[123] F. A. Ran, P. D. Hsu, J. Wright, V. Agarwala, D. A. Scott, and F. Zhang, “Genome engineering using the CRISPR-Cas9 system,” Nat. Protoc., vol. 8, no. 11, pp. 2281–2308, Nov. 2013, doi: 10.1038/nprot.2013.143.

[124] N. C. Stewart SA, Dykxhoorn DM, Palliser D, Mizuno H, Yu EY, An DS, Sabatini DM, Chen IS, Hahn WC, Sharp PA, Weinberg RA, “Lentivirus-delivered stable gene silencing by RNAi in primary cells,” RNA, vol. 9, no. 4, pp. 493–501, Apr. 2003, doi: 10.1261/rna.2192803.

[125] Y. Sancak et al., “The Rag GTPases Bind Raptor and Mediate Amino Acid Signaling to mTORC1,” Science (80-.)., vol. 320, no. 5882, pp. 1496–1501, Jun. 2008, doi: 10.1126/science.1157535.

[126] B. Wang, X, Seed, “A PCR primer bank for quantitative gene expression analysis,” Nucleic Acids Res., vol. 31, no. 24, pp. 154e – 154, Dec. 2003, doi: 10.1093/nar/gng154.

[127] A. Spandidos, X. Wang, H. Wang, and B. Seed, “PrimerBank: a resource of human and mouse PCR primer pairs for gene expression detection and quantification,” Nucleic Acids Res., vol. 38, no. suppl_1, pp. D792–D799, Jan. 2010, doi: 10.1093/nar/gkp1005.

